# Confidence Sets for Cohen’s *d* Effect Size Images

**DOI:** 10.1101/2020.05.21.108761

**Authors:** Alexander Bowring, Fabian Telschow, Armin Schwartzman, Thomas E. Nichols

## Abstract

Current statistical inference methods for task-fMRI suffer from two fundamental limitations. First, the focus is solely on detection of non-zero signal or signal change, a problem that is exasperated for large scale studies (e.g. UK Biobank, *N* = 40, 000+) where the ‘null hypothesis fallacy’ causes even trivial effects to be determined as significant. Second, for any sample size, widely used cluster inference methods only indicate regions where a null hypothesis can be rejected, without providing any notion of spatial uncertainty about the activation. In this work, we address these issues by developing spatial Confidence Sets (CSs) on clusters found in thresholded Cohen’s *d* effect size images. We produce an upper and lower CS to make confidence statements about brain regions where Cohen’s *d* effect sizes have exceeded and fallen short of a *non-zero* threshold, respectively. The CSs convey information about the magnitude and reliability of effect sizes that is usually given separately in a *t*-statistic and effect estimate map. We expand the theory developed in our previous work on CSs for %BOLD change effect maps (Bowring et al., 2019) using recent results from the bootstrapping literature. By assessing the empirical coverage with 2D and 3D Monte Carlo simulations resembling fMRI data, we find our method is accurate in sample sizes as low as *N* = 60. We compute Cohen’s *d* CSs for the Human Connectome Project working memory task-fMRI data, illustrating the brain regions with a reliable Cohen’s *d* response for a given threshold. By comparing the CSs with results obtained from a traditional statistical voxelwise inference, we highlight the improvement in activation localization that can be gained with the Confidence Sets.

## 1. Introduction

Online dating has transformed the love-seeking game forever. Whereas romantic partners would historically first encounter each other face-to-face, brought together by a mutual friend or family member, in recent times these rituals of connection have been largely replaced by social networks and matchmaking websites (Rosenfeld et al., 2019). While it was perhaps inevitable that technologies of the Digital Age would take a hold on our pursuit to find a partner, what may be more surprising is the influence the internet has had on the final outcomes of a marriage itself. In an investigation analyzing survey data from over 19,000 married American respondents, it was reported that virtual dating avenues may have helped to improve the prospects of finding a long and happy relationship (Cacioppo et al., 2013). With overwhelming statistical evidence, the results of this study found that spouses who had met their partner online were more likely to be satisfied with their marriage (p < 0.001) and less likely to divorce (*p* < 0.002).

Under closer inspection, however, these results are not all that they may seem. After this research was published, it was pointed out that the actual sizes of the observed effects were tiny (Nuzzo, 2013, 2014). Specifically, although the data had shown higher levels of marriage happiness for couples who met online compared to offline, the difference in means was from 5.5 to 5.6 on a 7-point scale; in terms of divorce rates, the deviation between groups worked out as one more break-up for every 100 marriages.

In this case study, the outcomes of the experiment were misconstrued due to an infamous pitfall with statistical testing known as the ‘fallacy of the null hypothesis’ (Rozeboom, 1960). The problem stems from the fact that statistical models conventionally assume mean-zero noise, when in reality all sources of noise will never cancel. Consequentially, the smallest of effects will always become statistically significant given a sufficiently large sample size (Meehl, 1967), even if they have little practical value or are unlikely to be replicable across repeated analyses (Button et al., 2013).

This issue has become topical within the functional magnetic resonance imaging (fMRI) community due to the arrival of population-scale neuroimaging datasets. While fMRI has tradition-Preprint submitted to Journal Name May 21, 2020 ally been a ‘small *N* ‘ enterprise, with typical sample sizes of 20 to 30 subjects (Poldrack et al., 2017), datasets such as the Human Connectome Project (HCP, *N* = 1, 200) and UK Biobank (*N* = 40, 000+) are now giving researchers the opportunity to analyze data acquired from tens of thousands of participants. These projects promise to transform our understanding of brain function, and are already yielding rich results (Miller et al., 2016, David C. Van Essen, 2016). However, in this setting the standard mass-univariate approach to functional brain data analysis has become obsolete: with ample power to detect all effects, statistical analysis of high quality fMRI data has been shown to lead to almost universal brain activation, even when stringent thresholding methods are applied (Gonzalez-Castillo et al., 2012).

In a more general analysis context, there are still a number of limitations with traditional fMRI inference techniques. Currently, the most popular method for overcoming the multiple comparisons problem is to threshold the statistical results with cluster-extent based thresholding (Carp, 2012), involving a two-step procedure: first a primary voxelwise threshold is applied to the statistic map, usually in correspondence with an uncorrected significance level (e.g. *α* = 0.001), creating clusters of voxels whose statistic values have all surpassed the threshold. Then, in order to control the family-wise error (FWE) rate, a cluster-extent threshold *k* is determined based on the distribution of cluster sizes obtained under the null-hypothesis of no activation, and the final results are computed as all suprathreshold clusters with a spatial extent larger than *k*.

As the FWE-corrected *p*-value is determined by cluster size, one of the main drawbacks of this procedure is that the significance of specific voxels can not be determined, and the most we can assert is that activation has occured *somewhere* inside a given cluster (Woo et al., 2014). It is therefore impossible to pinpoint the precise source of the activation when a cluster covers multiple anatomical regions, and the spatial specificity of the inference diminishes the larger a cluster becomes. Another problem with this approach is that no information is provided in regards to the spatial variation of significant clusters. For instance, if a single fMRI study was repeated many times using different groups of participants, there would be variation in the sizes and shapes of the final activation clusters, yet current statistical results have no way to convey this variability.

In our previous work (Bowring, Telschow, Schwartzman, and Nichols (2019) (*BTSN*)), we helped to address these issues by developing Confidence Sets (CSs) for inference on %BOLD change effect size maps. Unlike traditional hypothesis testing methods, where inference is only provided in terms of the presence of an effect, the CSs made simultaneous confidence statements about the precise brain regions where raw effect sizes had exceeded, and fallen short of, a *non-zero* %BOLD threshold. Here, we set out to adapt the CSs for application to standardized Cohen’s *d* maps (i.e. %BOLD change divided by population standard deviation) that are more commonly used to provide effect size estimates complementing the statistical results obtained from an fMRI one-sample *t*-test. For a cluster-forming threshold *c* and a predetermined confidence level 1 – α, the Cohen’s *d* CSs comprise of two sets: the upper CS (denoted 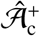), containing all voxels declared to have a true Cohen’s *d* effect size greater than *c;* and the lower CS 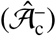, for which all voxels *outside* this set are declared to have a true Cohen’s *d* effect size less than *c*. The upper CS is smaller and nested inside the lower CS, and the assertion is made with (1 – *α*)100% confidence holding simultaneously for both regions. Note that *t*-tests, or any test statistics, are *not* suitable for building CSs, as they do not estimate a population quantity and become arbitrarily large for increasing sample sizes.

The statistical characteristics of Cohen’s *d* effect size maps are fundamentally different to the raw %BOLD images that motivated the CSs in our previous work. Our main contributions with this effort are modifications to the methods used in *BTSN* to create procedures for obtaining Cohen’s *d* CSs on fMRI data with desirable finite-sample performance. In particular, we apply recent results from the bootstrapping literature and a variance-stabilizing transformation method to ultimately propose three separate algorithms for computing Cohen’s *d* CSs. The first algorithm is motivated by asymptotic properties of the Cohen’s *d* sampling distribution, and provides a framework for the two remaining methods which employ further adjustments to optimize the finite-sample performance of the CSs. We assess the performance off all three methods on a range of simulated synthetic 2D and 3D signals representative of fMRI clusters, and find that the two latter procedures are effective even when the sample size is modest (*N* = 60). Finally, we apply the three procedures to Human Connectome Project working memory task data, operating on Cohen’s *d* effect maps, where we obtain CSs for a variety of cluster forming thresholds. By comparing the CSs with results obtained from a traditional statistical voxelwise inference, we highlight the improvement in activation localization that can be provided with the Confidence Sets.

The remainder of this manuscript is organized as follows: First, we describe the problem of obtaining Confidence Sets for Cohen’s *d* images, exemplifying the key differences which distinguish Cohen’s *d* from the %BOLD effect size. We then derive properties of the Cohen’s *d* estimator, before adapting the methods developed in *BTSN* to propose three separate algorithms to compute Cohen’s *d* CSs. We assess the empirical coverage performance of each of these methods on 2D and 3D Monte Carlo simulations, and finally, present the CSs obtained from applying each algorithm to Human Connectome Project working memory task-fMRI dataset.

## 2. Theory

### 2.1. From %BOLD to Cohen’s d

For a compact domain *S* ⊂ ℝ^*D*^, e.g. *D* = 3, for *i* = 1, …, *N* consider the one-sample model at location ***s*** ∈ *S*,

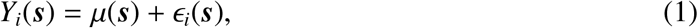

where *Y*_1_(***s***),…, *Y_N_*(***s***) are the observations at ***s***, *μ*(***s***) is the true underlying mean intensity across the observations, and *∈*_1_(***s***),…, *∈_N_*(***s***) are i.i.d. mean-zero errors with common variance *σ*^2^(***s***) and some unspecified spatial correlation. We are motivated by the setting of a group-level task-fMRI analysis, where *μ*(***s***) represents the true mean %BOLD change across the group, and each observation *Y_i_*(***s***) is the %BOLD response estimate map obtained by applying a first-level model to the *i*th participant’s functional data. (Note, while we focus on the one-sample model here, the method may also generalize for application to the general linear model ***Y***(***s***) = ***Xβ***(***s***) + *ϵ*(***s***). See the end of Section 5.1 for more details.)

We wish to make inference on the Cohen’s *d* effect size, defined as the true mean %BOLD change divided by the population standard deviation,

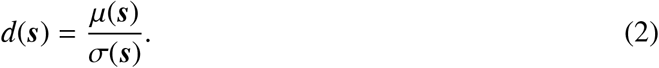

Specifically, we are interested in the brain regions where *d*(***s***) has exceeded, and fallen short of, a fixed threshold *c*, indicated by the noise-free, population cluster defined as:

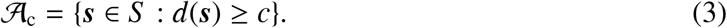

Since 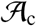 is unknown, we pursue a method for constructing pairs of spatial CSs: an upper set 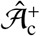 and a lower set 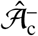, that we are confident surround the true excursion set 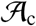 (i.e. 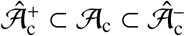) for a desired confidence level of, for example, 1 – *α* = 95%. Such a method lets us assert with 95% confidence that all voxels *contained* in the upper CS 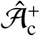 have a Cohen’s *d* effect size *greater* than, for example, *c* = 0.8, and simultaneously, we are 95% confident all voxels *outside* the lower CS 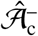 have a Cohen’s *d* effect size *less* than 0.8. Here, we emphasize the classic frequentist connotation of the term ‘confidence’; letting 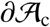 denote the boundary of 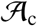, then precisely, there is a probability of 1 – *α* that the region 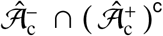 computed from a future experiment fully encompasses the true set boundary 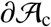. In this sense, the set difference of the upper and lower CS, 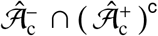, is similar to a standard confidence interval.

In *BTSN*, we adapted the mathematical theory first proposed in Sommerfeld, Sain, and Schwartzman (2018) (SSS) to obtain CSs for inference on the mean %BOLD change effect *μ*(***s***). Let 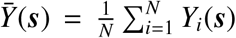, the sample mean %BOLD change. Then subject to continuity of the relevant fields and some basic conditions on the error terms *ϵ_i_*(***s***), for the excursion set 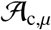 of voxels with a true %BOLD effect size greater than *c*,

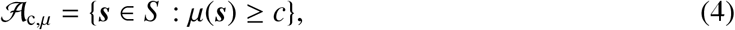

we showed that for a critical constant *k*, the upper and lower CSs constructed as

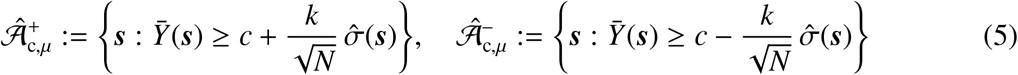

give asymptotic nominal coverage for enveloping the true 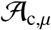 in terms of the mean %BOLD change effect size. Further to this, we proposed a Wild *t*-Bootstrap method for determining the critical value *k*, and demonstrated that on applying this method the CSs were also valid for data with smaller sample sizes.

We now seek to develop a similar methodology for the Cohen’s *d* effect size. However, the statistical properties of the Cohen’s *d* estimator 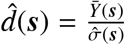 are considerably different to the sample mean 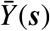. To provide a visual intuition of this in the case of Gaussian data, in Figure 1 we display images of both of these fields from a 2D simulation over a square region *S* = 100 × 100. For *N* = 60 subjects, we simulated a toy run of the signal-plus-noise model in (1) where the true underlying signal *μ*(***s***) was a linear ramp effect increasing from a magnitude of 0 to 10 in the *x*-direction while remaining constant in the *y*-direction (Figure 1(a)). To the signal we added subject-specific Gaussian noise *ϵ_i_*(***s***), smoothed with a 3 voxel FWHM Gaussian kernel and then re-normalized to have a spatially constant standard deviation of *σ(****s***) = 1. Notably, in this setup the true Cohen’s *d* field *d*(***s***) was identical to *μ*(***s***).

**Figure 1:**
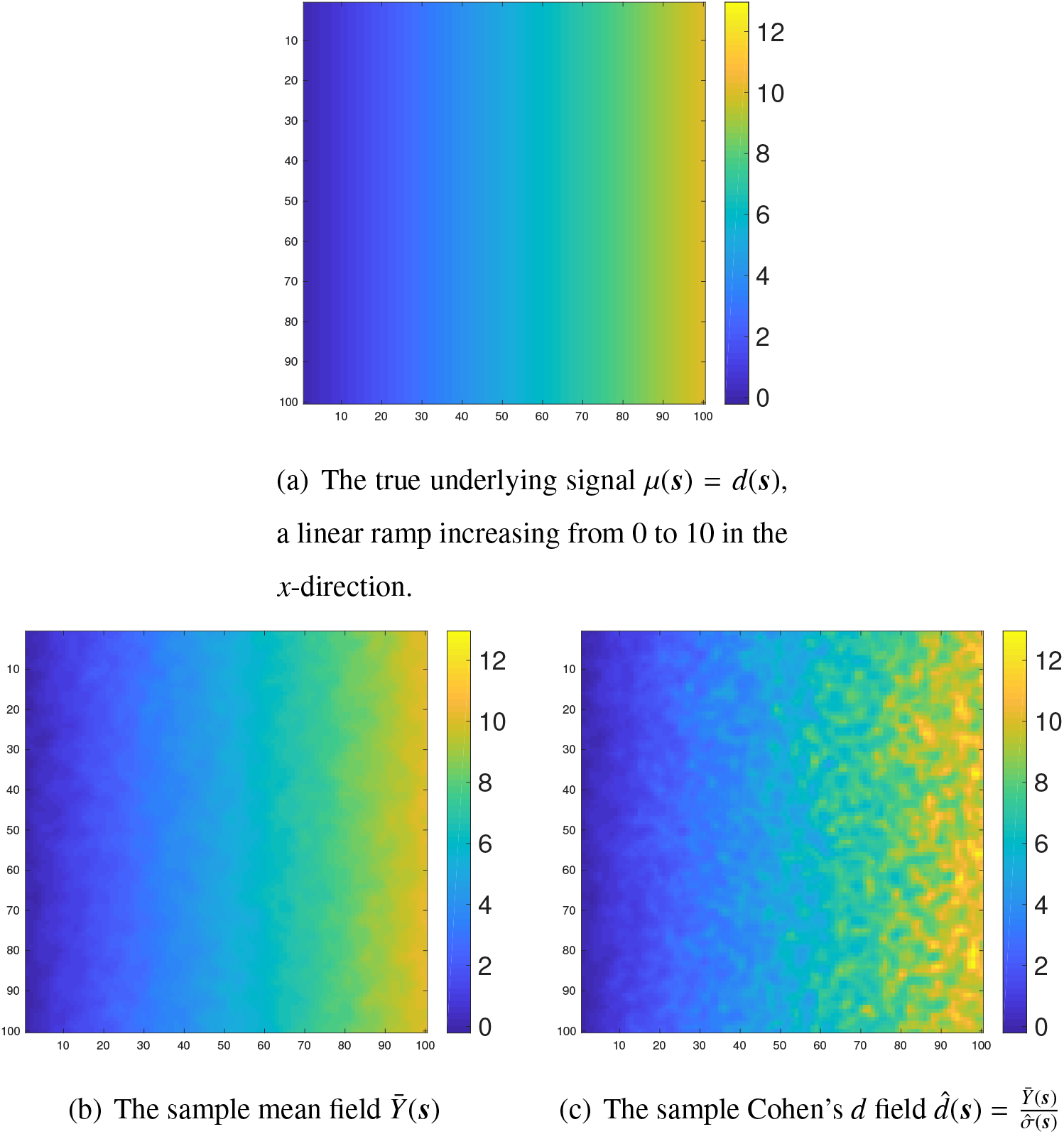
Visualizing the differences between the sample mean and sample Cohen’s *d* field. For *N* = 60 subjects, we simulated a signal-plus-noise model where the true underlying mean signal *μ*(***s***) was a linear ramp increasing from 0 to 10 across the region (a). To each subject we added Gaussian noise with a homogeneous variance, so that the true Cohen’s *d* effect *d*(**s**) was equal to the group mean signal *μ*(***s***). While the sample mean image 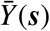 is uniformly smooth across the region (b), the sample Cohen’s *d* field 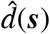 becomes rougher from left to right (c).

In Figure 1(b) and Figure 1(c) we show the sample mean and sample Cohen’s *d* fields from this simulation. While the sample mean image is uniformly smooth across the space, the Cohen’s *d* field becomes more speckled, i.e. more variable, for increasing true *d*(***s***). In the following sections we will show that the sample Cohen’s *d* is a biased estimator of the true underlying effect size, and that the sample variance of Cohen’s *d* changes systematically with *d*(***s***), before proposing our theoretical adjustments to the methods presented in *BTSN* to obtain CSs for Cohen’s *d* effect size images.

### 2.2. Limiting Properties of the Cohen’s d Estimator

Motivated by the example in Fig. 1, we now consider the one-sample model given in (1) with the additional assumption that the error fields are Gaussian. In this case, the data are i.i.d. 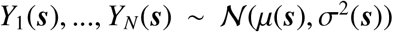 and the error terms *ϵ*_1_(***s***), …,*ϵ_N_*(***s***) are i.i.d. from a mean zero Gaussian random field *ϵ*(***s***) such that for all ***s, t*** ∈ *S*,

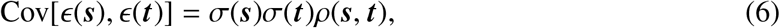

where *ρ*(***s, t***) denotes the population correlation coefficient between points ***s*** and ***t*** in the error field. The sample mean and sample standard deviation for this model are defined as

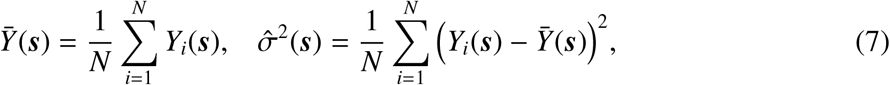

respectively.

We wish to understand the limiting structure of the Cohen’s *d* estimator 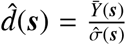. Applying the multivariate central limit theorem to the sample moments at ***s*** and ***t*** yields the asymptotic joint distribution:

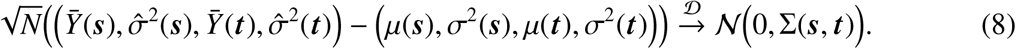

where the covariance matrix Σ(***s, t***) is given by

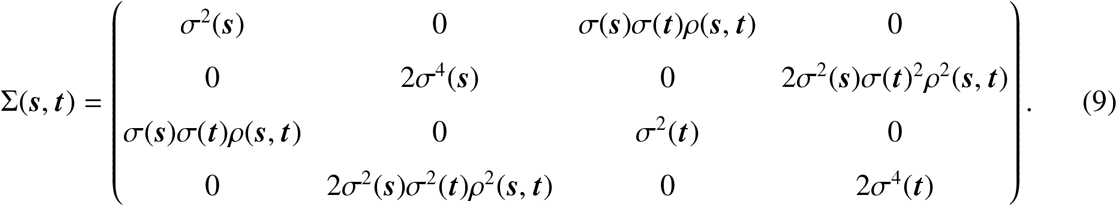

For the function 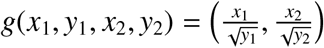, application of the delta method yields

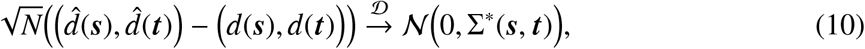

where

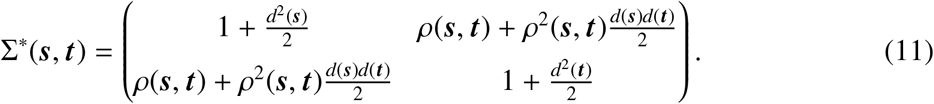

Therefore, the limiting field of the Cohen’s *d* estimator 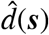 is asymptotically normal with asymptotic variance 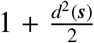. As alluded to in the previous section, it is notable that unlike the sample mean, the asymptotic variance and spatial correlation of the Cohen’s *d* estimator are dependent on the underlying true effect size. In the upcoming section, we will use these properties to motivate a construction for Cohen’s *d* Confidence Sets.

### 2.3. Confidence Sets for Cohen’s d Effect Size Images

Once again, consider the model outlined at the start of Section 2.1. For clarity, we reiterate that the spatial CSs for the raw %BOLD change field *μ*(***s***) of focus in our previous work took the form of (5), where *k* was determined via a Wild *t*-Bootstrap procedure. This construction of the CSs was motivated by the limiting properties of the field

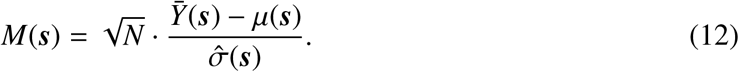

In particular, letting 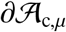 denote the boundary of 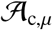 defined in (4), then on a neighbourhood *U* of 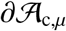, it is assumed in SSS that *M*(***s***) converges weakly to a smooth Gaussian field *G*(***s***) on *U* with mean zero, unit variance, and with the same (unknown) spatial correlation as each of the *ϵ_i_*.

In the previous section, for the Gaussian one-sample model we derived the convergence in distribution of the function

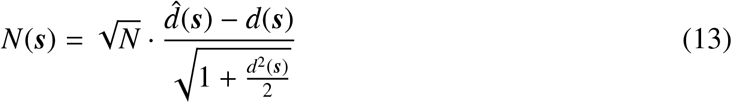

to a Gaussian field 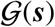 with mean zero, unit variance, and covariance structure

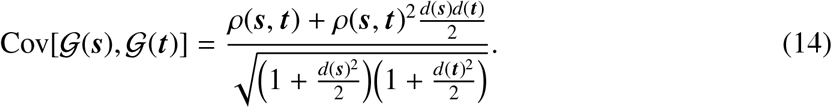

This suggests a natural analog to the construction of CSs in (5) for the Cohen’s *d* effect size given by

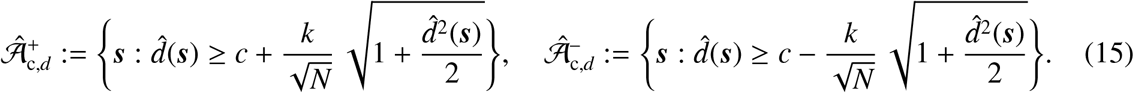

Ideally, we wishto apply the same Wild *t*-Bootstrap procedure described in Section 2.2 of *BTSN* to approximate the limiting field 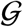 in order to determine *k*. However, we will now show that such an approach is not viable for Cohen’s *d*, before proposing a modified procedure to solve the problem. Going forward our focus will primarily be on the Cohen’s *d* effect size, and thus for brevity, we will drop the subscript from our notation and refer to the Cohen’s *d* CSs above as 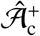 and 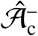 respectively.

### 2.4. Modified Residuals for the Cohen’s d Wild t-bootstrap

In *SSS*, it was shown that the limiting coverage of the CSs for the %BOLD effect size *μ*(***s***) is governed by the maximum distribution of the limiting Gaussian field *G*(***s***) on the boundary 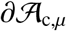, such that

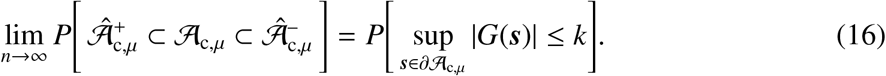

Since the limiting Gaussian field *G*(***s***) is unknown, in *BTSN* we implemented a Wild *t*-Bootstrap procedure to approximate *G*(***s***) on the boundary 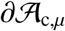. Defining the standardized residuals,

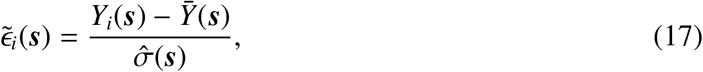

the Wild *t*-Bootstrap approximating field is given by

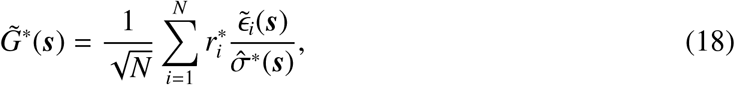

where the 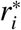 are i.i.d. Rademacher variables (i.e. each 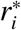 takes the value of-i ori with probability 1/2), and 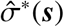 is the standard deviation of the current realization of bootstrapped residuals 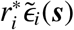. The asterisk (*) indicates that 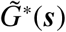 is one of many bootstrap samples; in practice, we would obtain a large number *B* of bootstrap samples 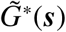, and approximate *k* as the (1 – *α*)100 percentile of the *B* suprema 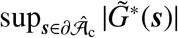.

While this method is valid in regards to %BOLD change, for Cohen’s *d* we demonstrate that asymptotically the correlation structure of the approximating field 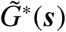 is incorrect. Consider again the Gaussian model in Section 2.2. In this instance, the covariance of the approximating field is:

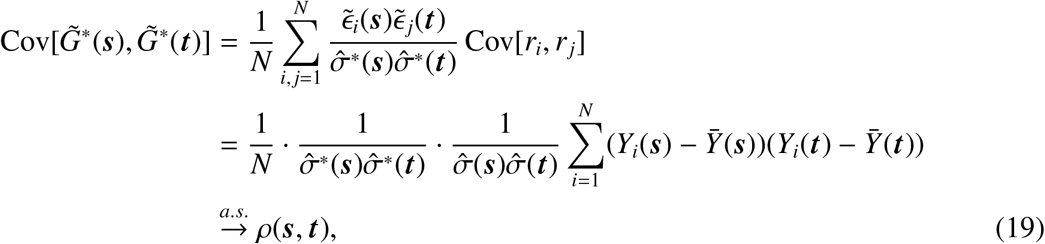

where we note that since the standardized residuals 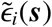 are asymptotically Gaussian with unit variance, the bootstrap estimate of the standard deviation of the standardized residuals 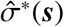 converges almost surely to 1. Since (19) and (14) don’t agree except for the complete null case of *d*(***s***) = 0 everywhere, we conclude that the covariance of the approximating field does not converge to the true covariance of the limiting field 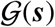.

To solve this problem, we implement a Taylor expansion transformation recently proposed in Telschow, Davenport, and Schwartzman (2020) to construct modified residuals with the desired limiting properties. Motivated by the delta method procedures used in Section 2.2, an estimation of the residual field for a single subject *i* is given by:

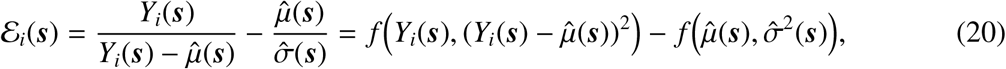

where 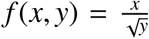. A first-order Taylor expansion of *f* (*x, y*) about the point 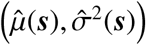 yields the approximating Cohen’s *d* residuals:

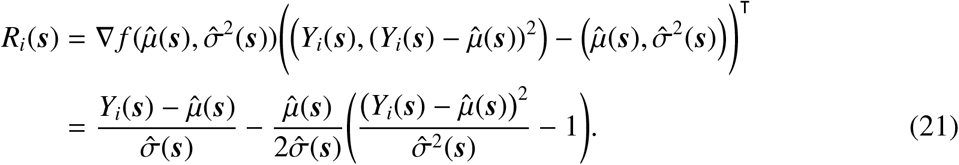

Normalizing by the estimated standard deviation of the limiting field 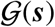, we obtain the modified standardized residuals:

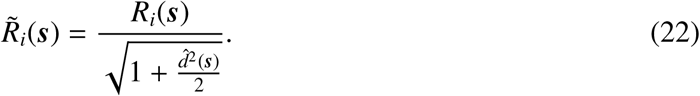

In Telschow et al., it is shown that the limiting covariance of 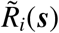 is equal to the covariance function of 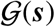. Therefore, a modification of (18) leads us to the Cohen’s *d* version of the Wild *t*-Bootstrap approximating field,

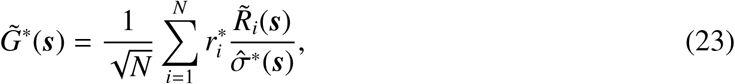

where now 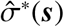 is the standard deviation of the bootstrapped Cohen’s *d* residuals 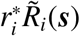.

While normalization of the *R_i_*(***s***) by an estimator of the standard deviation of 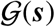 provides us with residuals that have the correct limiting properties, for application to fMRI data we wish to optimize the bootstrap in smaller sample sizes. In this regard, it may be preferable to standardize the *R_i_*(***s***) using an estimator tailored to the sample. Noting that the sample mean of the approximating residuals, 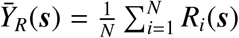, is equal to zero for all *N*, letting

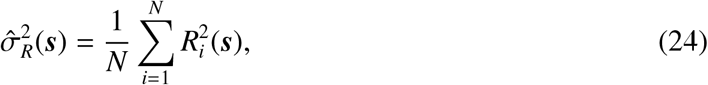

then an alternative to (22) is to normalize the Cohen’s *d* residuals by their sample standard deviation so that

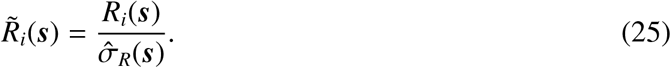

These standardized residuals can then be used for the Wild *t*-Bootstrap approximating field given in (23). In this case, the sample standard deviation should also be accounted for in the formation of the CSs, suggesting an alternate construction to (15) given by

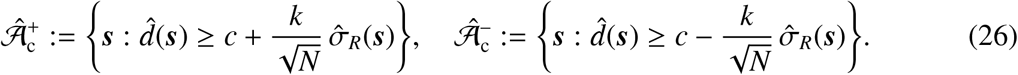

In Section 3.1, we assess the performance of the CSs on synthetic data when the residuals are standardized using either the estimated limiting variance 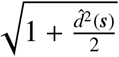 or the sample standard deviation 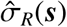.

### 2.5. Finite Properties of the Cohen’s d Estimator and a Variance-stabilizing Transformation

Up to now, we have motivated two possible constructions for Cohen’s *d* CSs ((15) and (26)) using the limiting properties of the Cohen’s *d* estimator. Here, we will draw our attention to the distributional properties of 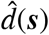 for finite samples to make further improvements on these methods, and introduce another novel procedure for obtaining CSs based on Gaussianizing the distibution of 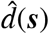.

Again, assuming the Gaussian model described in Section 2.2, observing that the Cohen’s *d* estimator can be expressed in the form,

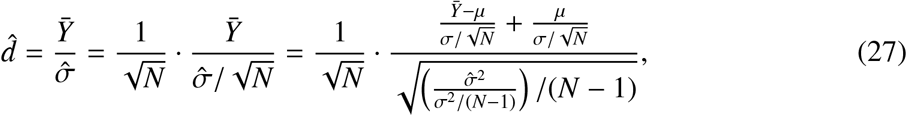

from the RHS of the equality we deduce that 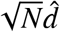 is characterized by a noncentral *t*-distribution with noncentrality parameter 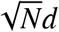 and *N* – 1 degrees of freedom.

Letting

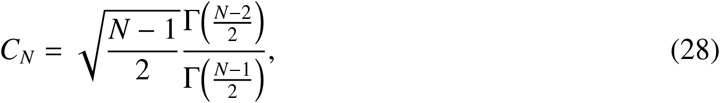

where Γ denotes the gamma function, then the expectation of 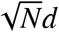 is given by

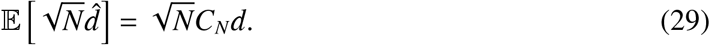

Therefore, unlike the sample mean, the Cohen’s *d* estimator is biased. To improve the performance of the CSs for small sample sizes, we will account for this bias in the formulation of the CSs. A well-known approximation of *C_N_* is

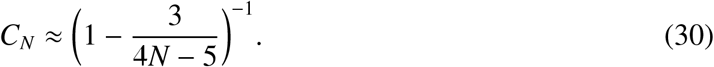

Therefore, a bias-corrected version of the CS construction in (15) given by

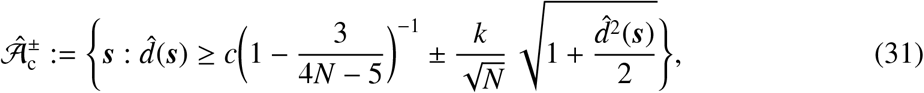

and similarly, a bias-corrected version of the alternate construction in (26) given by

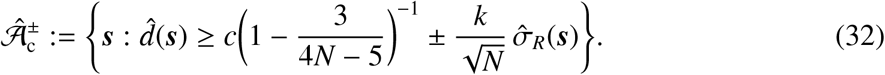

In addition to the formation of the CSs, for any application to real data, the Wild *t*-Bootstrap described in Section 2.4 must be applied over an approximation of the boundary 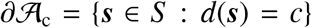. Taking into consideration the bias of the Cohen’s *d* estimator, we will use the plug-in boundary:

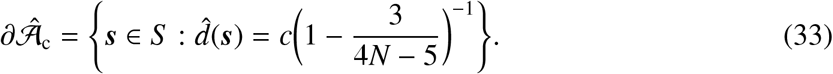

The noncentral *t*-distribution is asymmetric unless *μ* = 0; in general, the size of the asymmetry scales with the magnitude of the noncentrality parameter and is inversely proportional to the degrees of freedom. Therefore, we expect the distribution of the Cohen’s *d* estimator to be highly skewed when the true effect size is large and the sample size is small. This conflicts with the symmetric construction of the upper and lower CSs 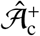 and 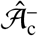 given in (31) and (32), suggesting that the coverage performance of these two methods may decline in such situations.

To account for skewness, we adapt a method originally proposed in Laubscher (1960) to stabilize the variance of the noncentral *t*, transforming to a distribution which is approximately Gaussian, and hence, symmetric. Letting

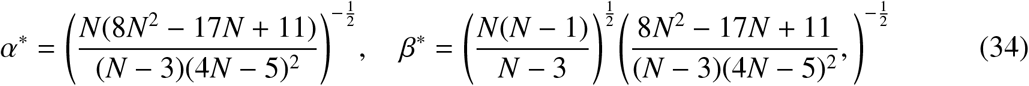

in Appendix A we show that the variance-stabilizing transformation of d is given by:

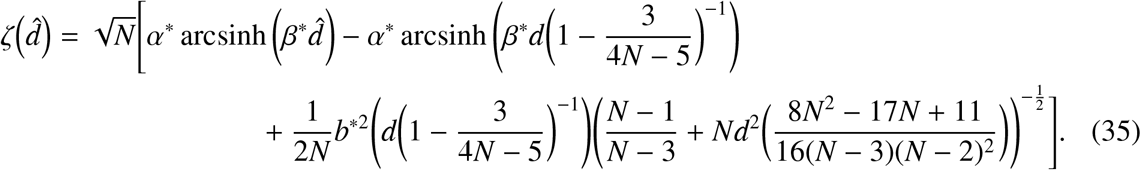

Numerical work presented in Laubscher shows that the 90th percentile value of 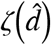 in (35) closely estimates *ϕ*^−1^(0.9) forarange oftrue effect sizes *d* when the sample size is larger than 40, suggesting that – for moderate sample sizes – the distribution of 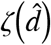 is approximately Gaussian.

By the monotonicity of the mapping *x* ↦ *α*^*^ arcsinh *β*^*^*x*, the variance-stabilizing transformation provides a further possibility for constructing the CSs in the transformed space 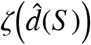. Reconstructing (35), the transformed CSs are given by:

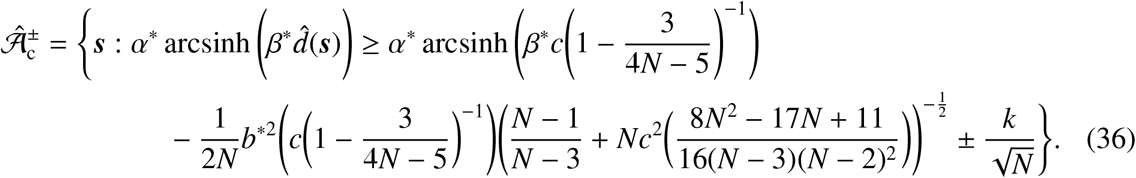

In this case, the Cohen’s *d* residuals given in (21) for the Wild *t*-Bootstrap must also be modified. An estimation of the transformed residual field for a single subject i is given by

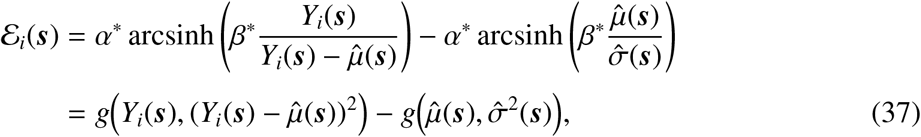

where the function 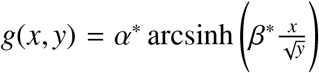. Similarly to the methods applied in Section 2.4, a first-order Taylor expansion of *g*(*x, y*) about the point 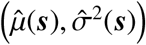 obtains the transformed Cohen’s *d* residuals

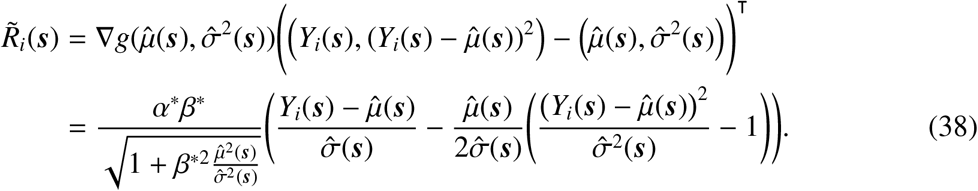

In practice, the critical value *k* in (36) is computed by applying the Wild *t*-Bootstrap over 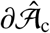 using the transformed Cohen’s *d* residuals given above for the bootstrap approximating field in (23).

For coherence, we will now formalize the complete procedures to obtain CSs for each of our three proposed CS constructions in (31), (32), and (36).

### 2.6. Three Algorithms for Computing Cohen’s d CSs

Based on our derivations up to this point, we give three algorithms to compute Cohen’s *d* CSs for data modelled within the one-sample model in Section 2.1. While the first two algorithms are similar, the key difference separating these methods is the estimator of the variance used in the formation of the CSs and for standardizing the Cohen’s *d* residuals. We first describe Algorithm 1., where we use 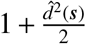 as the estimator of the variance, motivated by the variance of the limiting field 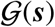 derived in Sections 2.2 and 2.3.

#### Algorithm 1.

*For observations Y*_1_(***s***), …, *Y_N_*(***s***) *modelled by the one-sample linear model in (1), the following procedure yields CSs for the Cohen’s *d* image* 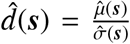 *corresponding to a fixed threshold c and confidence level* (1 – *α*)%.

1. *For each observation, Y_i_*(***s***), *let* 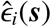 *denote the residual field*, 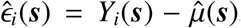. *Then compute the Cohen’s *d* residuals as*

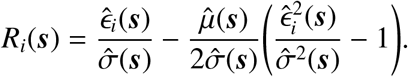
2. *Normalize the Cohen’s *d* residuals by the estimated limiting standard deviation of the Cohen’s *d* image to obtain the standardized residuals,*

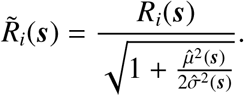
3. *Draw N i.i.d. Rademacher variables* 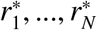, *and compute the Wild t-Bootstrap approximating field*,

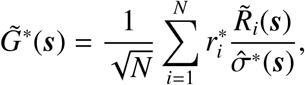

*where* 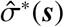 is the bootstrap standard deviation of the bootstrapped residuals 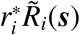.
4. *Obtain the value* 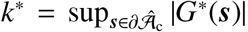, *using the bias-corrected estimator of the boundary* 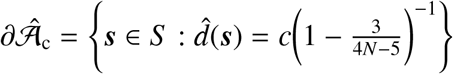
5. *For a large number of bootstrap replications B repeat steps 3. and 4., obtaining the empirical distribution of the absolute maximum* 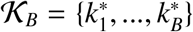. *Compute k as the* (1 – *α*) *percentile of* 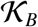.
6. *Obtain the Cohen’s *d* CSs*,

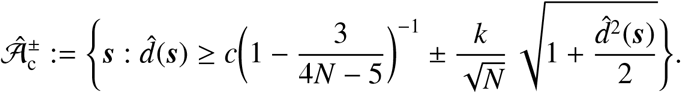

For Algorithm 2., we use the sample variance of the Cohen’s *d* residuals 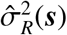 as the variance estimator, motivated by our workings in Section 2.4.

#### Algorithm 2.

*For observations Y*_1_(***s***),…,*Y_N_*(***s***) *modelled by the one-sample linear model in (1), the following procedure yields CSs for the Cohen’s d image* 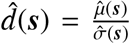 *corresponding to a fixed threshold c and confidence level* (1 – *α*)%.

1. *For each observation, Y_i_*(***s***), let 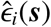 *denote the residual field*, 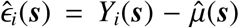. *Then compute the Cohen’s d residuals as*

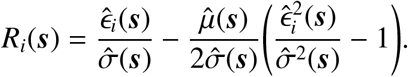
2. *Normalize the Cohen’s d residuals by their sample standard deviation to obtain the standardized residuals,*

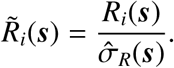
3. *Draw N i.i.d. Rademacher variables* 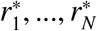, *and compute the Wild t-Bootstrap approximating field*,

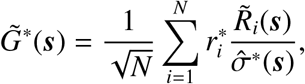

*where* 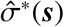 *is the bootstrap standard deviation of the bootstrapped residuals* 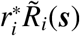.
4. *Obtain the value* 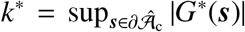, *using the bias-corrected estimator of the boundary* 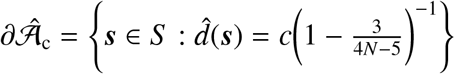.
5. *Fora large number of bootstrap replications B repeat steps 3. and 4., obtaining the empirical distribution of the absolute maximum* 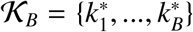. *Compute k as the* (1 – *α*) *percentile of* 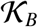.
6. *Obtain the Cohen’s d CSs*,

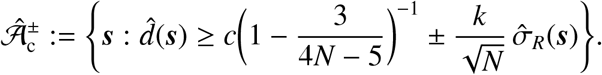

Finally, Algorithm 3 is based on the derivations in Section 2.5, transforming the estimated Cohen’s *d* image to a field which is approximately Gaussian. This is done to stabilize the variance and remove the skew of the Cohen’s *d* estimator, which may adversely effect the performance of the CSs. By the monotonicity of the transformation, the CSs obtained using this method are valid for inference on the true (un-transformed) Cohen’s *d* effect size.

#### Algorithm 3.

*For observations Y*_1_(***s***), …, *Y_N_*(***s***) *modelled by the one-sample linear model in (1), the following procedure yields CSs for the Cohen’s d image* 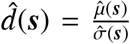 *corresponding to a fixed threshold c and confidence level* (1 – *α*)%.

1. *For each observation, Y_i_*(***s***), *let* 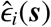 *denote the residual field*, 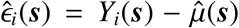. *Then compute the transformed, variance-stabilized Cohen’s d residuals as*

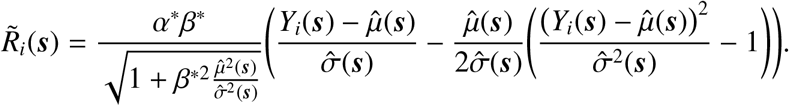
2. *Draw N i.i.d. Rademacher variables* 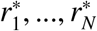, *and compute the Wild t-Bootstrap approximating field*,

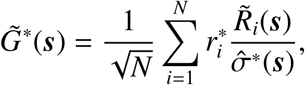

*where* 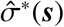 *is the bootstrap standard deviation of the bootstrapped residuals* 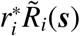.
3. *Obtain the value* 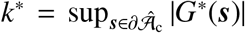, *using the bias-corrected estimator of the boundary* 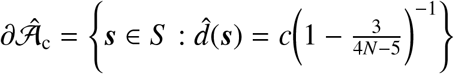
4. *Fora large number of bootstrap replications B repeat steps 3. and 4., obtaining the empirical distribution of the absolute maximum* 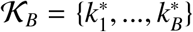. *Compute k as the* (1 – *α*) *percentile of* 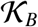.
5. *Obtain the Cohen’s d CSs*,

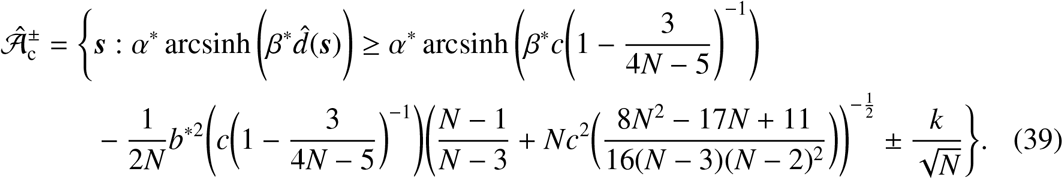

## 3. Methods

### 3.1. Simulation Setup

In this section we describe the settings used to evaluate the performance of each of the three algorithms for obtaining Cohen’s *d* CSs on synthetic data. For each method, we simulate 3000 independent samples of the Gaussian one-sample model

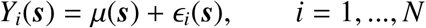

using a range of signals *μ*(***s***), Gaussian noise structures *ϵ_i_*(***s***) with stationary and non-stationary variance *σ*^2^(***s***), in two- and three-dimensional regions *S*. To compute the critical value *k*, we apply the given method’s Wild *t*-Bootstrap procedure with *B* = 5000 bootstrap samples on the estimated boundary 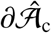 that must be used for application to real data. We obtain the boundary using the linear interpolation method described in Section 2.3 of *BTSN*. We then compute the empirical coverage using the interpolation assessment method described in Section 2.4 of *BTSN*, given as the percentage of trials for which the true thresholded signal 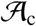 contains the upper CS 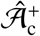 and is contained within the lowers CS 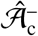 i.e. the proportion of trials for which 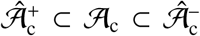. In each simulation, we apply the given method to sample sizes of *N* = 60, 120, 240 and 480, and for each of the three nominal coverage probability levels 1 – *α* = 0.80, 0.90 and 0.95.

### 3.2. 2D Simulations

We analyzed the performance of the three algorithms to obtain Cohen’s *d* CSs on a square region of size 100 × 100. For the true underlying signal *μ*(***s***) we considered two different raw effects: first, a linear ramp that increased from a magnitude of 0 to 1 in the *x*-direction while remaining constant in the *y*-direction. Second, a circular effect, created by placing a circular phantom of magnitude 1 and radius 30 in the centre of the search region, which was then smoothed using a3 voxel FWHM Gaussian kernel. If we were to assume that each voxel had a size of 2mm^3^, we note that this would amount to applying smoothing with a 6mm FWHM kernel, a fairly typical setting used in fMRI analyses.

To each of these signals we added subject-specific Gaussian noise *ϵ_i_*(***s***), obtained from smoothing white noise with a3 voxel FWHM Gaussian kernel, with homogeneous and non-homogeneous variance structures: the first noise field had a spatially constant standard deviation of 1, and therefore in this case the true Cohen’s *d* effect was identical to the underlying signal *μ*(***s***). The second field had a linearly increasing standard deviation structure in the *y*-direction from 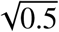 to 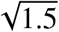 while remaining constant in the *x*-direction. Thus, the variance of this noise field spatially increased in the *y*-direction from 0.5 to 1.5 in a non-linear fashion.

The true Cohen’s *d* fields *d*(***s***) for the linear ramp signal with homogeneous and heterogeneous noise are shown in Figure 2. The corresponding Cohen’s *d* fields for the circular signal are shown in Figure 3. Altogether, for the three algorithms, the two underlying signals and two noise sources gave us 12 different simulation setups; for all of the simulations, we obtained Cohen’s *d* CSs for the noise-free cluster 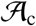 at a cluster-forming threshold of*c* = 0.8. In Chapter 2.2 of Cohen (2013), *d* = 0.8 was classified as a ‘large effect’; for group-level analyses of large-sample fMRI data with ample statistical power (such as the HCP or UK Biobank), effect sizes of this magnitude may be used to assess brain areas where practically significant activation has occurred.

**Figure 2:**
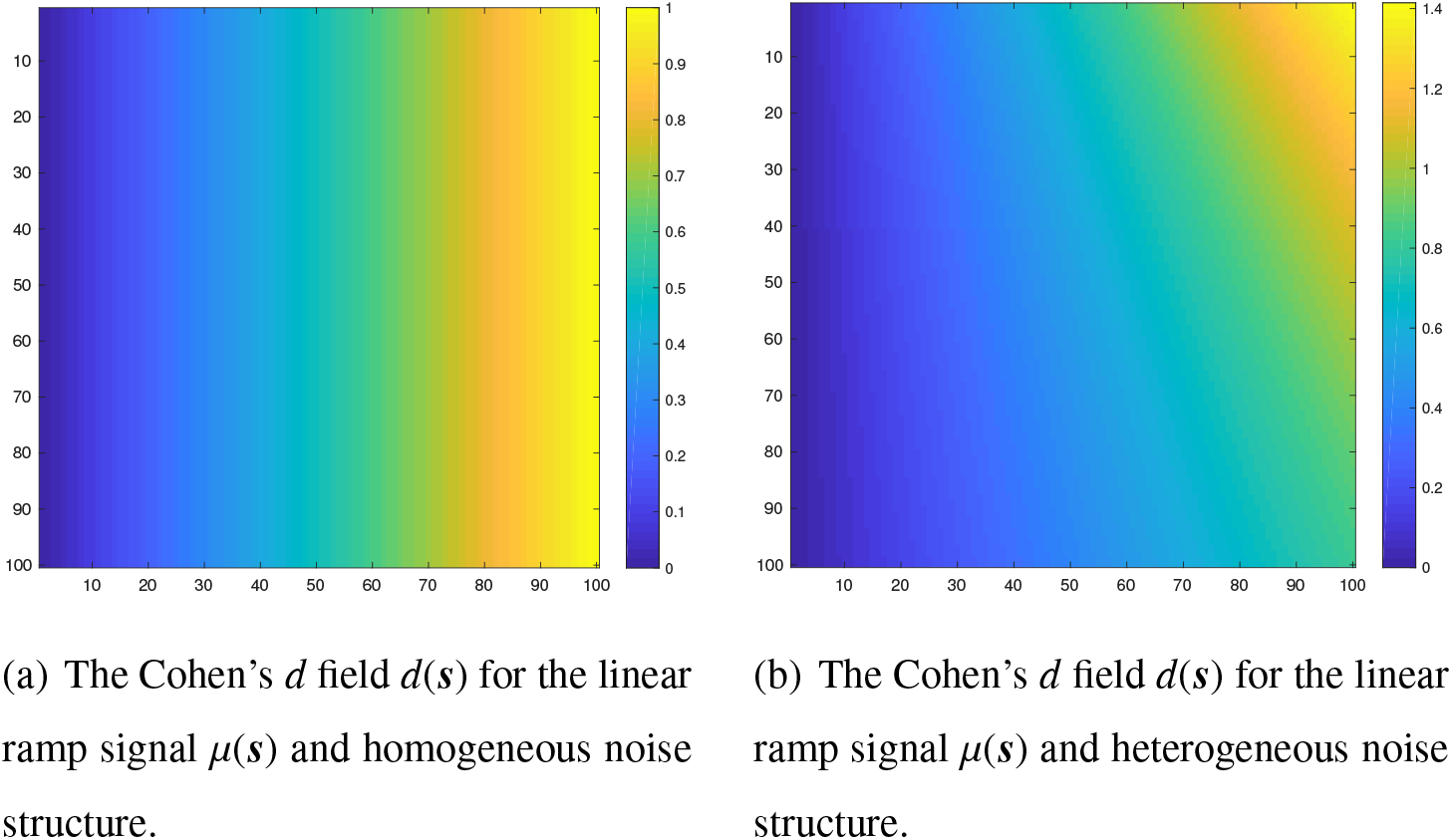
The two Cohen’s *d* effects corresponding to the linear ramp signal *μ*(***s***). On the left, the subject-specific Gaussian noise field *ϵ_i_*(***s***) has a spatially constant standard deviation of 1, and therefore *d*(***s***) = *μ*(***s***). On the right, *ϵ_i_*(***s***) had a spatially increasing standard deviation structure in the *y*-direction (from top-to-bottom), while remaining constant in the *x*-direction.

**Figure 3:**
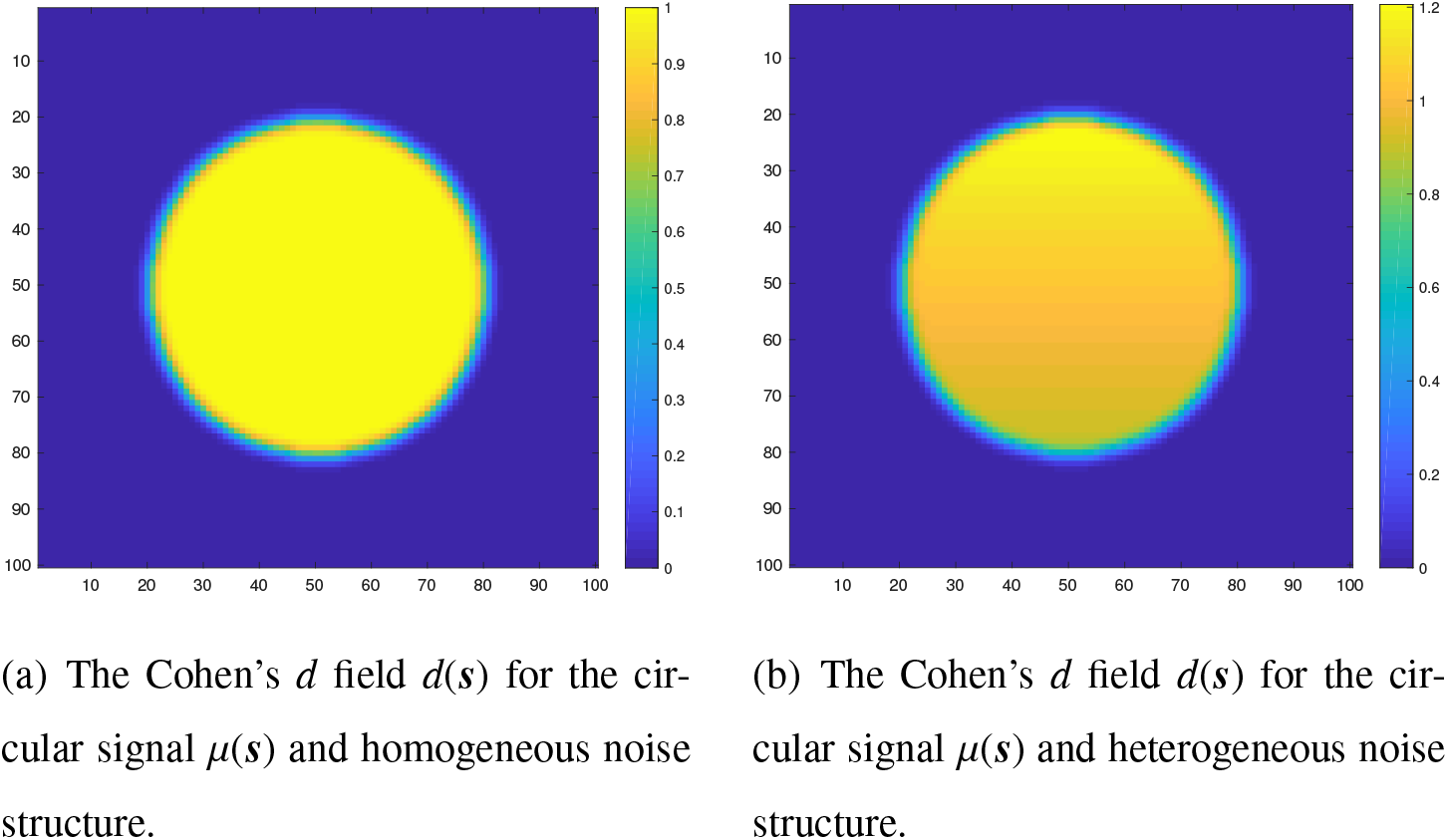
The two Cohen’s *d* effects corresponding to the circular signal *μ*(***s***). On the left, the subject-specific Gaussian noise field *ϵ_i_*(***s***) has a spatially constant standard deviation of 1, and therefore *d*(***s***) = *μ*(***s***). On the right, *ϵ_i_*(***s***) had a spatially increasing standard deviation structure in the *y*-direction (from top-to-bottom), while remaining constant in the *x*-direction.

### 3.3. 3D Simulations

Four signal types *μ*(***s***) were considered to analyze the performance of the three algorithms in three dimensions. The first three of these signals were generated synthetically on a cubic region of size 100 × 100 × 100: firstly, a small spherical effect, created by placing a spherical phantom of magnitude 1 and radius 5 in the centre of the search region, which was then smoothed using a 3 voxel FWHM Gaussian kernel. Secondly, a larger spherical effect, generated identically to the first effect with the exception that the spherical phantom had a radius of 30. Lastly, we created an effect by placing four spherical phantoms of magnitude 1 in the region of varying radii and then smoothing the entire image using a 3 voxel FWHM Gaussian.

Each of the images were rescaled after smoothing to have a maximum intensity of 1. For the small and large spherical effect an imagewise rescaling was applied, where all locations in the smoothed map were divided through by the maximum intensity across the region. For the final effect, because parts of the four spherical phantoms overlapped after smoothing, the signal intensities in these regions summed to greater than 1. In this case, we reduced the intensities in these areas to have a magnitude of 1 while leaving the rest of the image untouched. This ensured that the signal at the center of each spherical phantom had a magnitude of 1, coinciding with the previous small and large spherical effect signal types (see Fig. 4, the signal at the center of each spherical phantom in plot (c) is 1, in correspondence with the small and large spherical effects in plots (a) and (b)).

**Figure 4:**
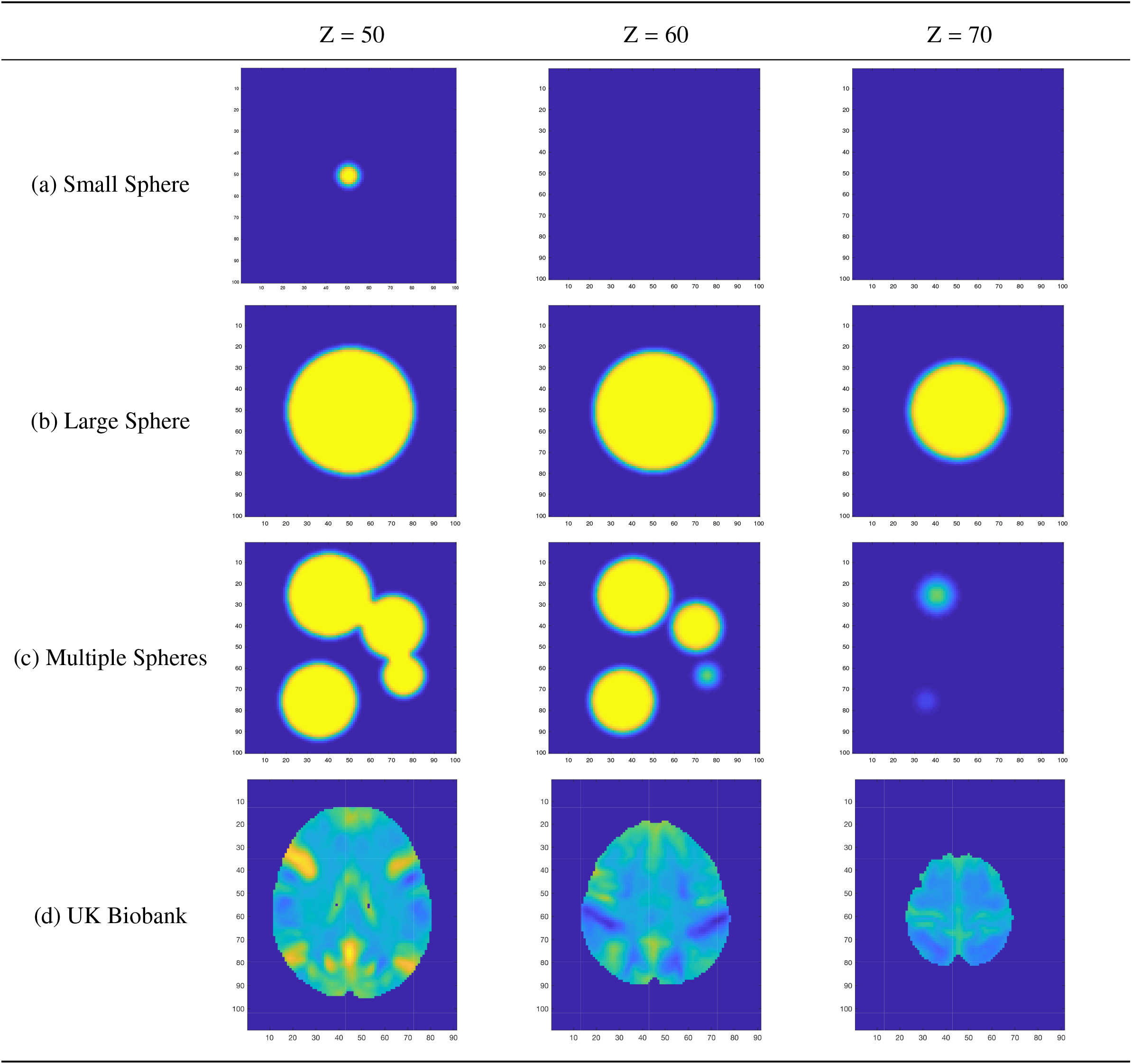
Four of the Cohen’s *d* fields *d*(**s**) used for the 3D simulations. Plots (a) to (c) show the Cohen’s *d* field for the three different spherical effects *μ*(***s***) when Gaussian noise with spatially homogeneous standard deviation was added to the signal. Plot (d) shows the Cohen’s *d* field corresponding to the UK Biobank full mean and standard deviation images. Note that the colormap limits for the first three Cohen’s *d* effect-size images are from 0 to 1, while the colormap limits for the UK Biobank image is from −0.8 to 0.8.

Similar to the two-dimensional simulations, for the three signals described above we added white noise smoothed using a 3-voxel FWHM Gaussian kernel with homogeneous and heterogeneous variance structures. The first noise field had a spatially constant standard deviation of 1, while the second field had a linearly increasing standard deviation in the *z*-direction from 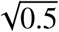 to 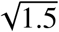, while remaining constant in both the x- and *y*-directions. As demonstrated for the 2D simulations in Figures 2 and 3, this lead to two different true Cohen’s *d* effect-size images *d*(***s***) corresponding to the homogeneous and heterogeneous standard deviation fields *σ*(***s***) used for the noise.

For the final signal type, we took advantage of big data that has been made available through the UK Biobank in an attempt to generate an effect that replicated the true %BOLD change induced during an fMRI task. We randomly selected 4000 subject-level contrast of parameter estimate result maps from the Hariri Faces/Shapes task-fMRI data collected as part of the UK Biobank brain imaging study. Full details on how the data were acquired and processed is given in *Miller et al. (2016), Alfaro-Almagro et al. (2018)* and the UK Biobank Showcase; information on the task paradigm is given in *Hariri et al. (2002)*. From these contrast maps, we computed a group-level full mean and full standard deviation image, considering all voxels where at least one subject had data (instead of discarding voxels with any missing data). In the final simulation, we used the group-level Biobank mean image as the true underlying signal *μ*(***s***) for each subject, and the full standard deviation image was used for the standard deviation of each simulated subject-specific Gaussian noise field *ϵ_i_*(***s***) added to the true signal. Because of the considerably large sample size of high-quality data from which these maps have been obtained, we anticipate that both of these images are highly representative of the true underlying fields that they approximate. Both images were masked using an intersection of all 4000 of the subject-level brain masks.

Once again, we smoothed the noise field using a 3 voxel FWHM Gaussian kernel; as the Biobank maps had voxel sizes of 2mm^3^, this equated to applying 6mm FWHM smoothing to the noise field of the original data.

In Figure 4, we show the true underlying Cohen’s *d* fields for the three synthetic 3D effects with homogeneous noise structure, and the Cohen’s *d* field corresponding to the UK Biobank full mean and standard deviation. For all four signal types, we obtained Cohen’s *d* Confidence Sets for the threshold *c* = 0.8.

### 3.4. Application to Human Connectome Project Data

To provide areal-data demonstration of the three methods proposed in this work, we computed Cohen’s *d* CSs on 80 participants data from the Unrelated 80 package released as part of the Human Connectome Project (HCP, S1200 Release) using all three algorithms described in Section 2.6. Cohen’s *d* CSs were obtained for the subject-level 2-back vs 0-back contrast maps from the working memory task results included with the HCP dataset. For a comparison with standard fMRI inference procedures, we also performed a traditional statistical group-level analysis on the data. A one-sample *t*-test was carried out in SPM, using a voxelwise FWE-corrected threshold of *p* < 0.05 obtained via permutation test with SPM’s SnPM toolbox.

For the working memory task participants were presented with pictures of places, tools, faces and body parts in a block design. The task consisted of two runs, where on each run a separate block was designated for each of the image categories, making four blocks in total. Within each run, for half of the blocks participants undertook a 2-back memory task, while for the other half a 0-back memory task was used. Eight EVs were included in the GLM for each combination of picture category and memory task (e.g. 2-back Place); we compute CSs on the subject-level contrast images for the 2-back vs 0-back contrast results that contrasted the four 2-back related EVs to the four 0-back EVs.

Imaging was conducted on a 3T Siemans Skyra scanner using a gradient-echo EPI sequence; TR = 720ms, TE = 33.1 ms, 208 × 180 mm FOV, 2.0 mm slice thickness, 72 slices, 2.0 mm isotropic voxels, and a multi-band acceleration factor of 8. Preprocessing of the subject-level data was carried out using tools from FSL and Freesurfer following the ‘fMRIVolume’ HCP Pipeline fully described in Glasser et al. (2013). To summarize, the fundamental steps carried out to each individual’s functional 4D time-series data were gradient unwarping, motion correction, EPI distortion correction, registration of the functional data to the anatomy, non-linear registration to MNI space (using FSL’s Non-linear Image Registration Tool, FNIRT), and global intensity normalization. Spatial smoothing was applied using a Gaussian kernel with a 4mm FWHM.

Modelling of the subject-level data was conducted with FSL’s FMRIB’s Improved Linear Model (FILM). The eight working task EVs were included in the GLM, with temporal derivatives terms added as confounds of no interest, and regressors were convolved using FSL’s default double-gamma hemodynamic response function. The functional data and GLM were temporally filtered with a high pass frequency cutoff point of 200s, and time series were prewhitened to remove autocorrelations from the data.

Mirroring the methods used in *BTSN*, we applied additional smoothing to the final contrast maps to mimic images smoothed using a 6mm FWHM Gaussian kernel. This is a more typical degree of smoothing applied to functional data than the 4mm kernel originally used in the HCP analysis pipeline.

## 4. Results

### 4.1. 2D Simulations

Empirical coverage results for each of the three algorithms are presented for the linear ramp signal in Figure 5 and for the circular signal in Figure 6. In both figures, on the top row we display the coverage results obtained when the standard deviation field of the noise was homogeneous across the region (corresponding to Fig. 2(a) for the linear ramp, Fig. 3(a) for the circle), and on the bottom row we display the equivalent results when the standard deviation field was spatially heterogeneous (Fig. 2(b) and Fig. 3(b) for the linear ramp and circle respectively).

**Figure 5:**
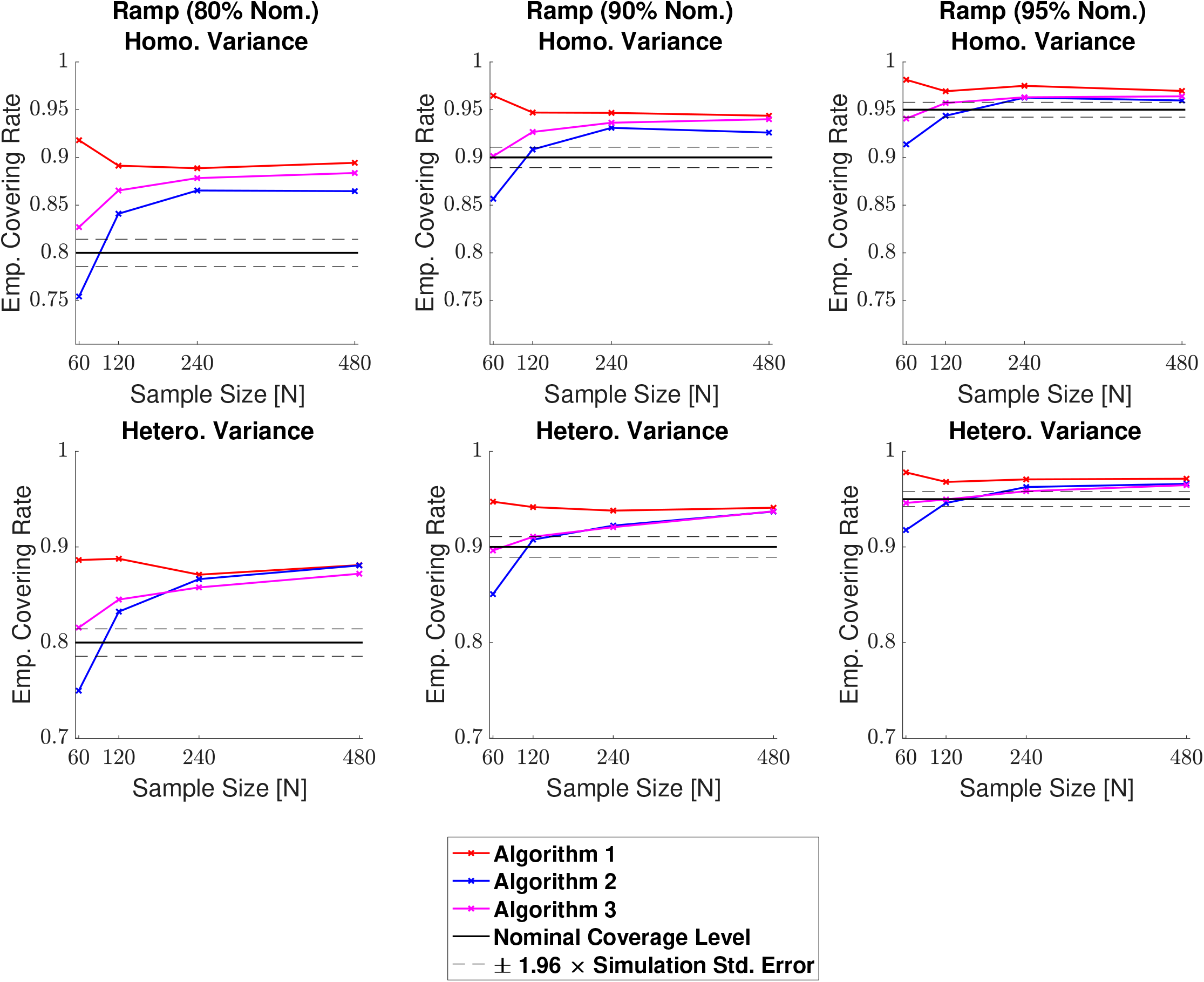
Coverage results for the linear ramp signal, with homogeneous (top row) and heterogeneous (bottom row) Gaussian noise structures. For large sample sizes the empirical coverage performance of all three algorithms was similar, hovering slightly above the nominal level in all simulations. For *N* = 60 the degree of over-coverage became larger for Algorithm 1., while empirical coverage for Algorithm 2. fell below the nominal target. Algorithm 3. performed best, with all results remaining particularly close to the nominal target level for simulations using a 95% confidence level (right plots).

**Figure 6:**
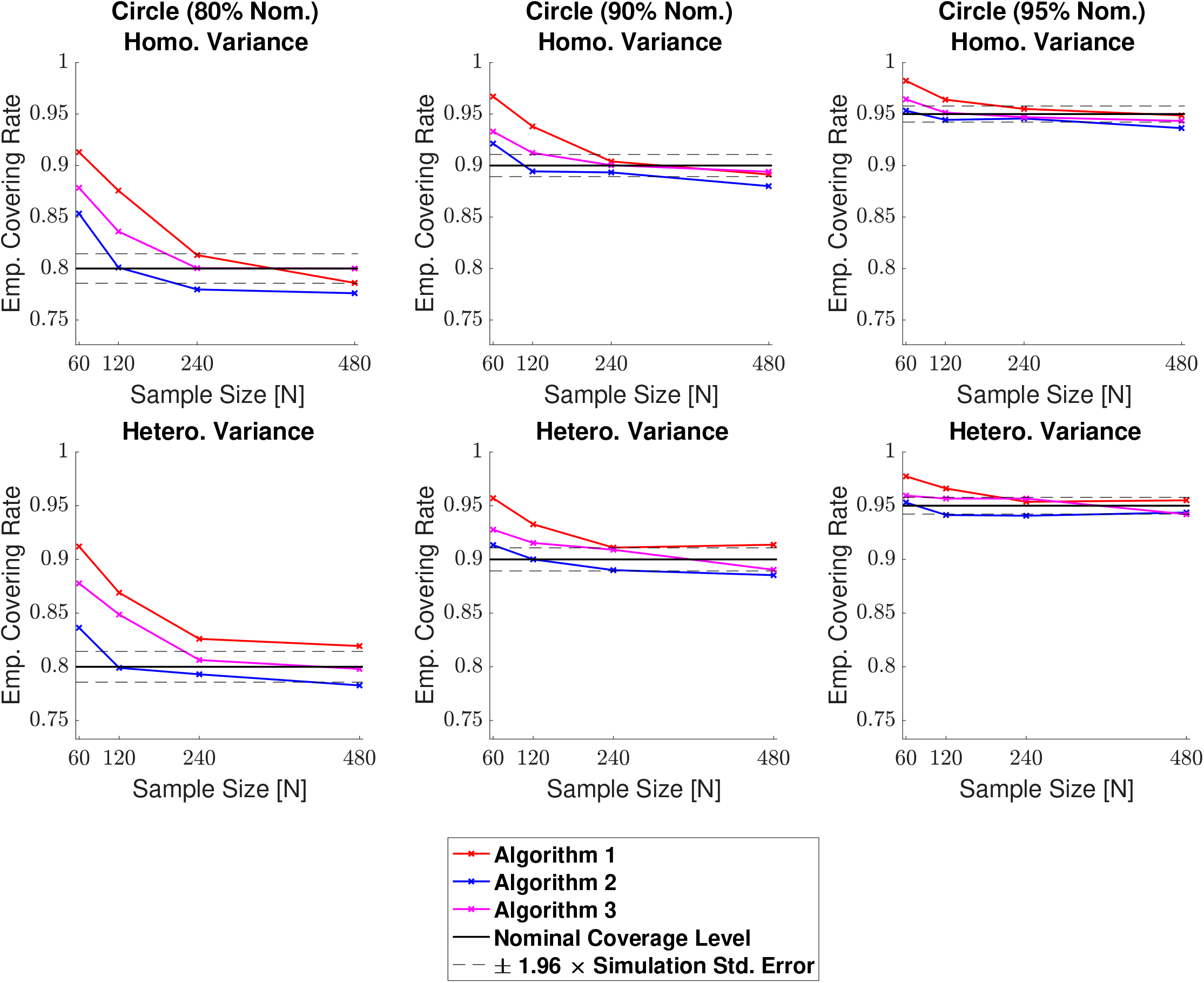
Coverage results for the circular signal, with homogeneous (top row) and heterogeneous (bottom row) Gaussian noise structures. All algorithms performed very well, and unlike the linear ramp, empirical coverage for all three methods converged quickly towards the nominal level. For smaller sample sizes there was a slight degree of over-coverage, most noticeably for simulations using the 80% nominal target. Overall, Algorithm 2. performed marginally better than the other two methods, and Algorithm 1. performed the worst.

For the linear ramp, across all confidence levels 1 – *α* = 0.80, 0.90, and 0.95 we generally observed valid over-coverage for all three algorithms, particularly when larger sample sizes were used. In all plots, it appears that the coverage rates for the three algorithms are converging to the same value, slightly above the nominal target. Specifically, for the nominal target level of 80%, in both the homogeneous and heterogeneous cases all empirical results seem to be converging to around 88% (Fig. 5, left-side plots). For the 95% target, the scale of disagreement between the empirical results and the nominal target is smaller; here, all coverage results hover close to 96% for *N* = 240 and 480 (Fig. 5, right-side plots).

While for larger sample sizes the performance of all three algorithms was similar, there was greater disparity between the methods for simulations using the small sample size of *N* = 60. Here, the empirical coverage results for Algorithm 1. were consistently higher than the other two methods, and at the same time, results for Algorithm 2. were always the smallest. Notably, unlike the other two methods, the empirical coverage rate for Algorithm 2. fell below the nominal level in all simulations with a sample size of *N* = 60.

For the circular signal, on the whole all three methods performed very well. In this instance, almost all empirical results for Algorithm 2. and Algorithm 3. lay within the 95% confidence interval of the nominal coverage rate (blue and magenta curves sandwiched between black dashed lines for all plots in Fig. 6), with Algorithm 2. performing marginally better. While we observed slight over-coverage with the three methods for *N* = 60, most substantially in simulations using the 80% nominal target (Fig. 6, left-side plots), empirical coverage converged towards the nominal level for all three algorithms.

Finally, the use of homogeneous or heterogeneous noise in the model had minimal impact on any of the algorithm’s empirical coverage performance for either of the signals. This is exemplified in Figs. 5 and 6, where in both cases the homogeneous coverage plots presented in the top row are almost identical to the corresponding heterogeneous plots shown below.

### 4.2. 3D Simulations

Empirical coverage results for each of the three algorithms are presented in Figures 7, 8, 9 and 10 respectively for each of the four 3D signal types displayed in Fig. 4 (small sphere, large sphere, multiple spheres, UK Biobank). For the spherical effects (Figs. 7, 8 and 9), on the top row we display the coverage results obtained when the standard deviation field of the noise was homogeneous across the region, and on the bottom row we display the equivalent results when the standard deviation field was spatially heterogeneous. For the UK Biobank signal (Fig. 10), the full standard deviation image computed from the UK Biobank data was used for the standard deviation field of the noise, and hence in this case there is only one row of results.

**Figure 7:**
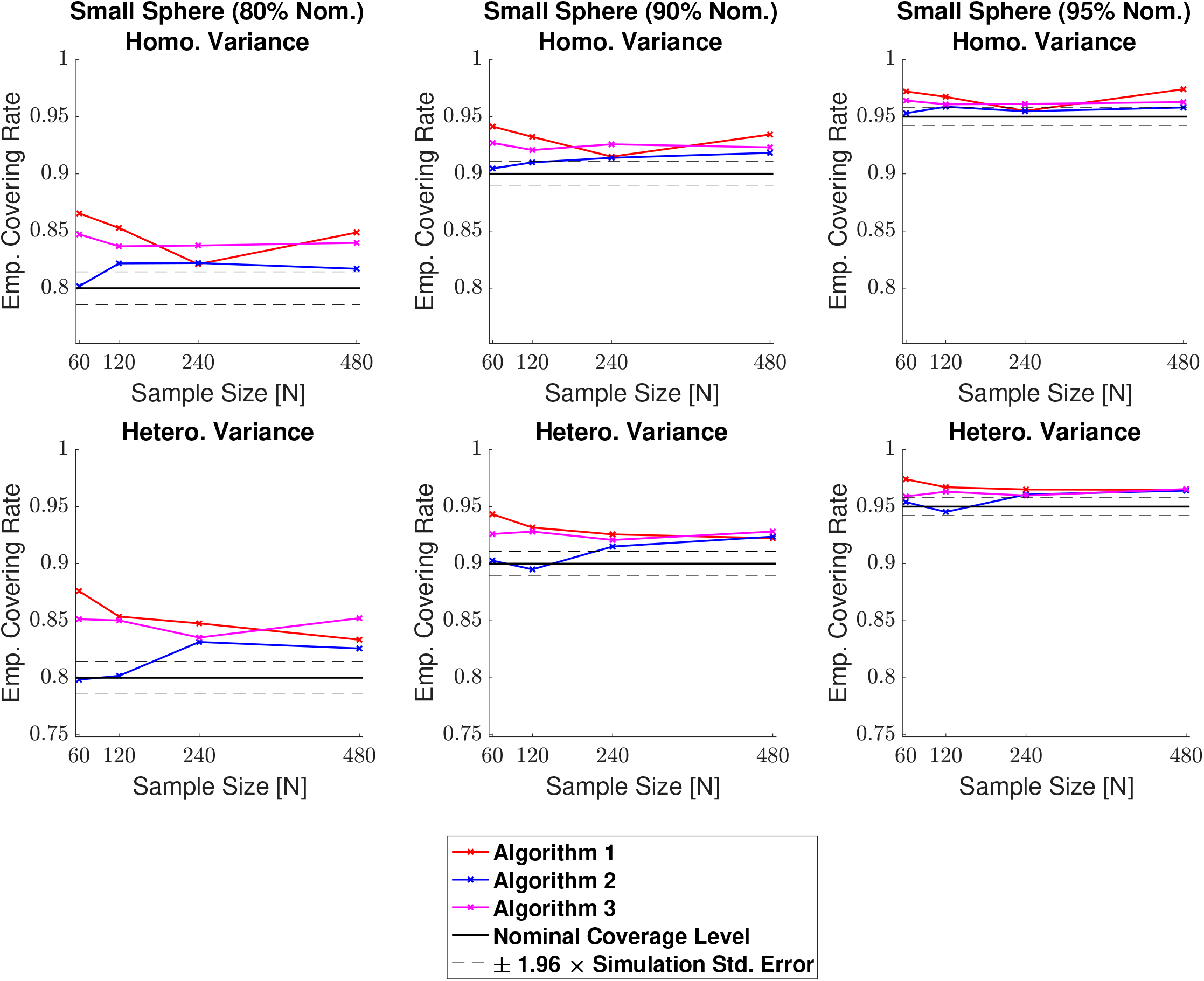
Coverage results for the small sphere signal type, with homogeneous (top row) and heterogeneous (bottom row) Gaussian noise structures. In general, empirical coverage remained above the nominal level across all simulations, and for the 95% confidence level (right plots), the results of all three methods fell close to the nominal target. All methods were robust as to whether the subject-level noise had homogeneous or heterogeneous variance structure. Because of this, there are minimal differences comparing the plots between both rows.

**Figure 8:**
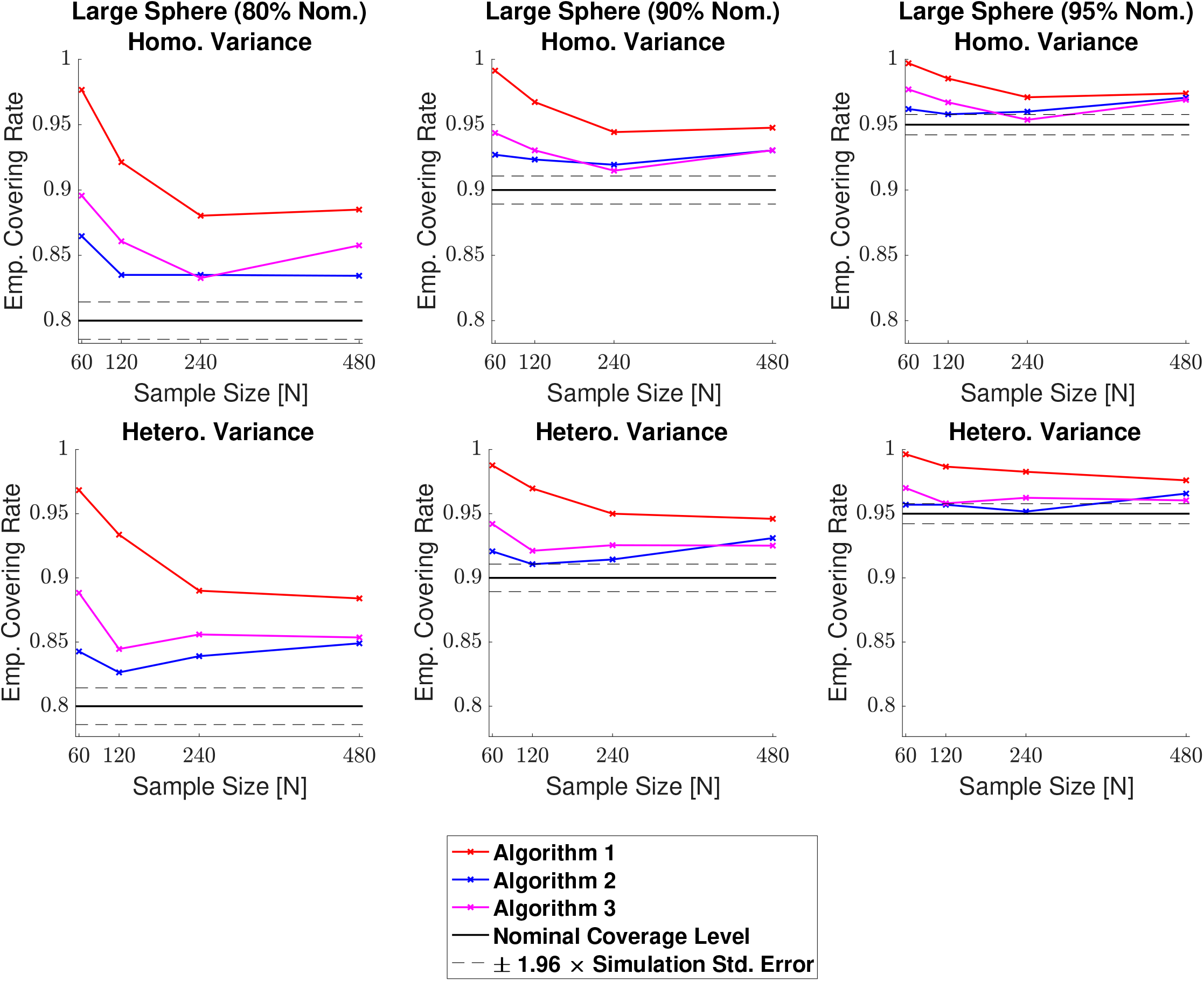
Coverage results for the large sphere signal type, with homogeneous (top row) and heterogeneous (bottom row) Gaussian noise structures. Compared with the small sphere results displayed in Fig. 8, empirical coverage results were higher for all three methods here. Algorithm 1. suffered from a particularly large degree of over-coverage for simulations with a small sample size. Coverage performance for Algorithm 2. and Algorithm 3. was closer in resemblance to the corresponding small sphere results, with Algorithm 2. performing slightly better. This suggests that both of these methods are fairly robust to changes in the boundary length.

**Figure 9:**
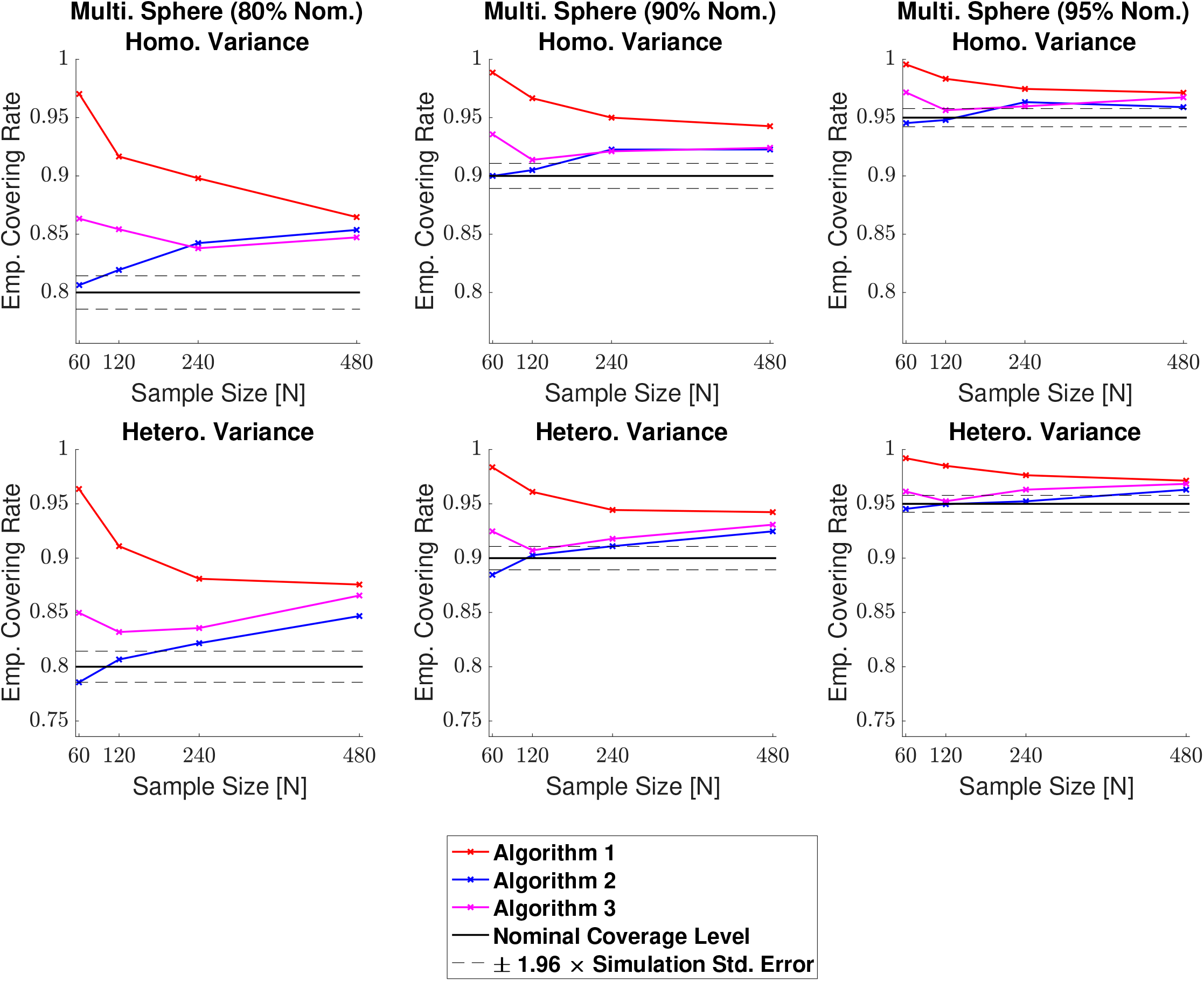
Coverage results for the multiple spheres signal type, with homogeneous (top row) and heterogeneous (bottom row) Gaussian noise structures. Algorithm 2. and Algorithm 3. both performed well, particularly for the 95% confidence level, where coverage levels remained in the vicinity of the 95% confidence interval of the nominal target. While Algorithm 2. was closer to the nominal target for *N* = 60, in some cases empirical coverage went slightly below the nominal level. In comparison to the other two methods, Algorithm 1. suffered from a large degree of over-coverage, which slightly improved as the sample size increased.

**Figure 10:**
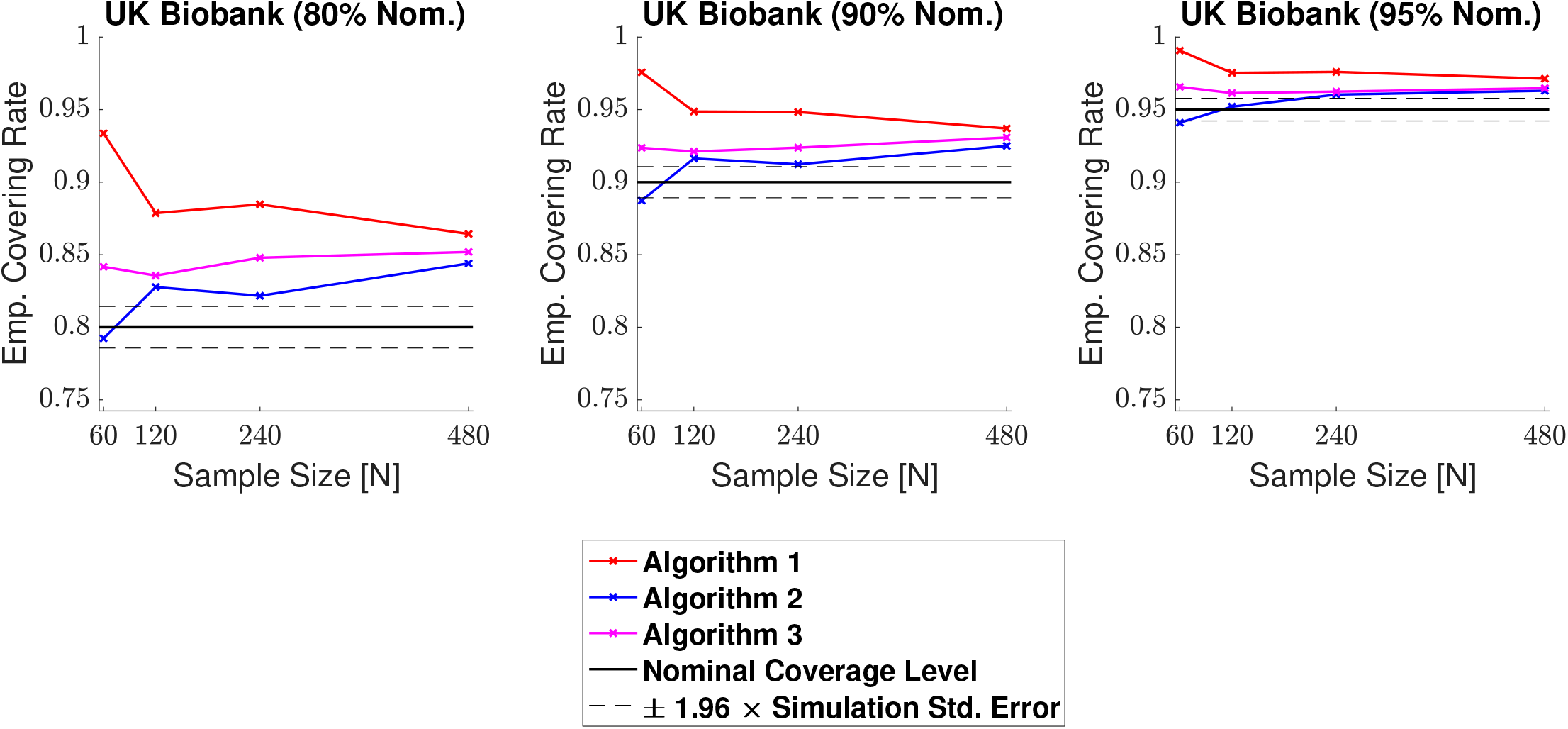
Coverage results for the UK Biobank signal type, where the full standard deviation image was used as the standard deviation of the subject-level noise fields. Coverage results here were similar to the results for the multiple spheres signal type shown in Fig. 9. Once again, both Algorithm 2. and Algorithm 3. performed well, with empirical coverage rates hovering above the nominal target for large sample sizes, while results for Algorithm 1. came further above the nominal level.

Across all 3D simulations, we observed consistencies between the results obtained with each of the three algorithms: in general, empirical coverage for all methods came above the nominal target, and similar to the 2D simulations, the extent of over-coverage was smaller when a larger confidence level was used. Comparing the three methods, coverage results for Algorithm 1. were considerably higher than the other two methods, particularly when a small sample size and confidence level were used. For the ‘large sphere’ and ‘multiple spheres’ signal types, Algorithm 1. suffered with over-coverage of above 15% in simulations with a sample of size of *N* = 60 and a nominal target level of80% (Figs. 8 and 9, left-side plots). For both of these signals, there was still a considerable amount of over-coverage when larger sample sizes of *N* = 1 20, 240 and (to a lesser degree) 480 were used. On the other hand, Algorithm 2. and Algorithm 3. performed similarly in large sample sizes across all simulations, with empirical coverage results coming slightly above the nominal target. Notably, both of these algorithms performed very well for simulations with a 95% nominal target level (all figures, right-side plots). Differences between these two methods were more distinguished for smaller sample sizes of *N* = 60 and 120, where coverage results for Algorithm 2. were moderately less than Algorithm 3. Consequentially, Algorithm 2.‘s results came closer to the nominal target here, although for the ‘multiple sphere’ and ‘UK Biobank’ signal types, in some cases Algorithm 2.‘s results fell *below* the nominal level (Figs. 9 and 10). Overall, empirical coverage for Algorithm 3. was the most uniform of the three methods with respect to changes in sample size.

Comparing Figs. 7 and 8, we observed a slight deterioration in the performance of all three algorithms when moving from the small sphere signal type to the large sphere. In particular, results obtained from applying the three methods to the large sphere fell further above the nominal target relative to the small sphere. This was most severe for Algorithm 1., where differences between the two sets of results were larger than 10% for the 80% confidence level (Figs. 7 and 8, left-side plots). These differences were comparatively marginal for Algorithm 2. and Algorithm 3., where we observed only a slight increase in empirical coverage. This would suggest that both of these methods are fairly robust to changes in the boundary length.

Finally, the use of homogeneous or heterogeneous noise in the model once again had very little impact on the performance of all three algorithms. Nevertheless, for simulations with small sample sizes, a heterogeneous noise structure led to a slight decrease in the empirical coverage results for Algorithm 2. and Algorithm 3. (Figs. 7, 8 and 9, left-side plots).

### 4.3. Human Connectome Project

Cohen’s *d* Confidence Sets obtained by applying Algorithm 3. to 80 subjects’ contrast data from the Human Connectome Project are shown in Figure 11. CSs computed on the same data using Algorithm 1. and Algorithm 2. are displayed in Figures S.1 and S.2 respectively. For each figure, we display the CSs obtained from applying the specified algorithm with three separate thresholds, *c* = 0.5, 0.8, and 1.2. These three Cohen’s *d* effect sizes were classified as medium, large, and very large in Cohen (2013).

**Figure 11:**
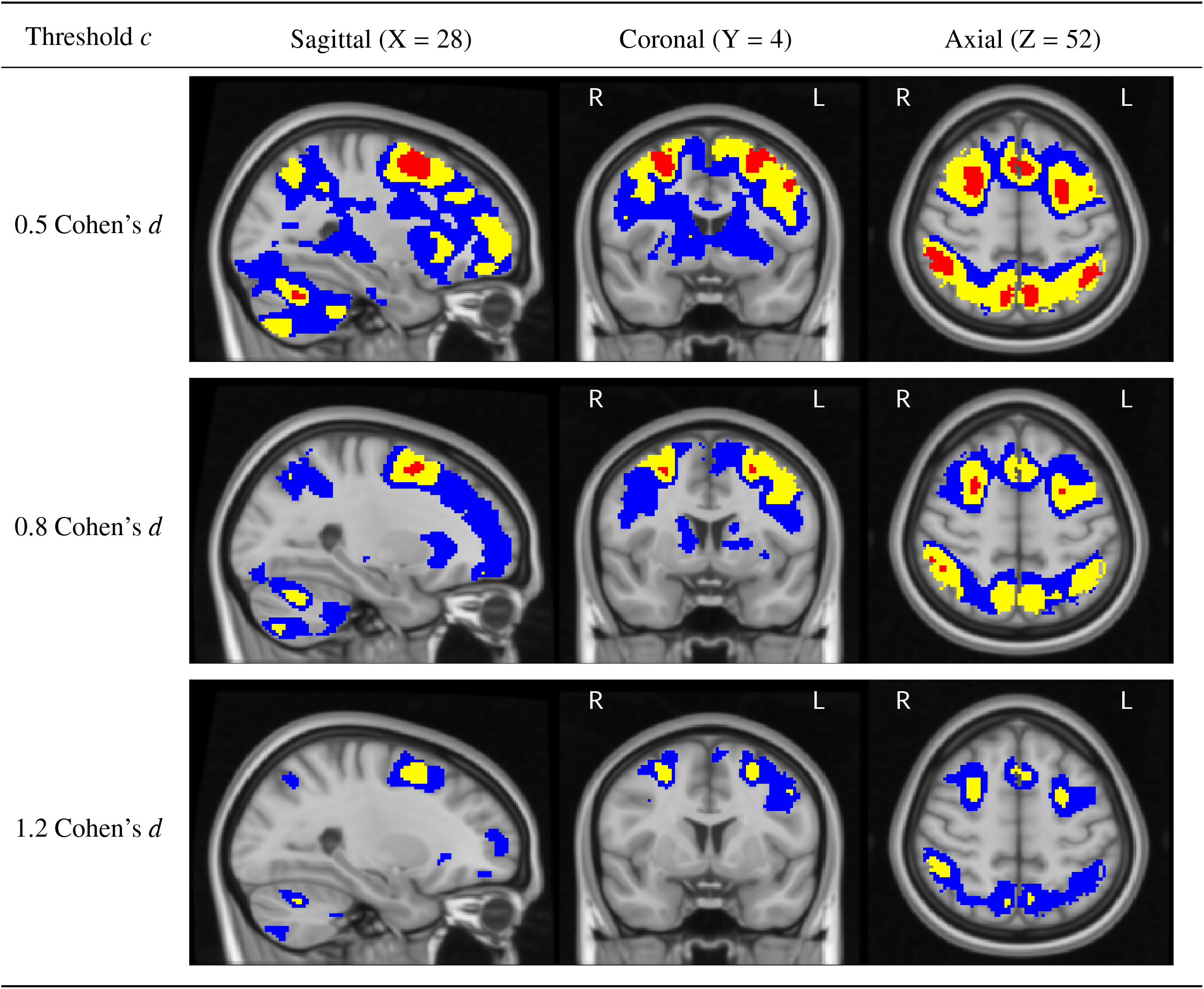
Slices views of the Cohen’s *d* Confidence Sets obtained from applying Algorithm 3. to the HCP working memory task data, using three Cohen’s *d* effect size thresholds, *c* = 0.5, 0.8 and 1.2. The upper CS 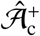 is displayed in red, and the lower CS 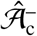 in blue. Yellow voxels represent the point estimate set 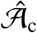, the best guess from the data of voxels that have surpassed the Cohen’s *d* threshold. The red upper CS has localized regions in the frontal gyrus, paracingulate gyrus, angular gyrus, cerebellum and precuneus which we can assert with 95% confidence have attained (at least) a 0.5 Cohen’s *d* effect size.

In the top plot of Fig. 11, the red upper CSs localized brain regions within the frontal cortex that are commonly associated with working memory. This included areas of the superior frontal gyrus (left and right, all slices), middle frontal gyrus (left, coronal slice), paracingulate gyrus (left and right, axial slice) and insular cortex. Other brain areas encapsulated inside the upper CS were the angular gyrus (left and right, axial slice), cerebellum (left and right, sagittal slice) and precuneus (left and right, axial slice). For all these regions, the method identified clusters of voxels where we can assert with 95% confidence there was a Cohen’s *d* effect size greater than 0.5.

By increasing the threshold to *c* = 0.8 (Fig. 11, middle plot), there was a shrinking of both the blue lower CSs and red upper CSs. Therefore, while we can confidently declare a medium effect size in all of the brain areas identified above, the quantity of voxels within each region that we can proclaim to have a large effect size is considerably smaller. In the case of the right cerebellar hemisphere (left, sagittal slice) and insular cortex, the upper CS vanished completely, indicating that the method did not locate any voxels in these regions where we can assert a Cohen’s *d* effect size greater than 0.8.

For the largest threshold assessed, *c* = 1.2, the red upper CS was empty, and hence we can not assert any region of the brain to to have attained a very large effect size. Notably, the yellow point estimate set contains a small but appreciable number of voxels, signifying that based on the data alone, these voxels were estimated to have a Cohen’s *d* effect size greater than 1.2. Conversely, the large quantity of (grey backgroud) voxels lying outside the blue lower CS in Fig. 11 imply an effect size *less* than 1.2 across the vast majority of the brain.

In Figure 12, the red upper CSs computed with Algorithm 3. are compared with the thresholded *t*-statistic map (green-yellow voxels) obtained from applying a one-sample *t*-test group-analysis to the 80 subjects’ contrast data, using a voxelwise FWE-corrected threshold of *p* < 0.05. This figure demonstrates the improved spatial specificity that can be provided with the CSs in comparison with the traditional approach. Specifically, while the thresholded statistic map contains one large cluster covering a sizeable portion of the parietal lobe across both brain hemispheres, the red upper CSs pinpoint precise areas in the precuneus and angular gyrus where a practically significant medium (or large) Cohen’s *d* effect size can be inferred (Fig. 12, axial slices).

**Figure 12:**
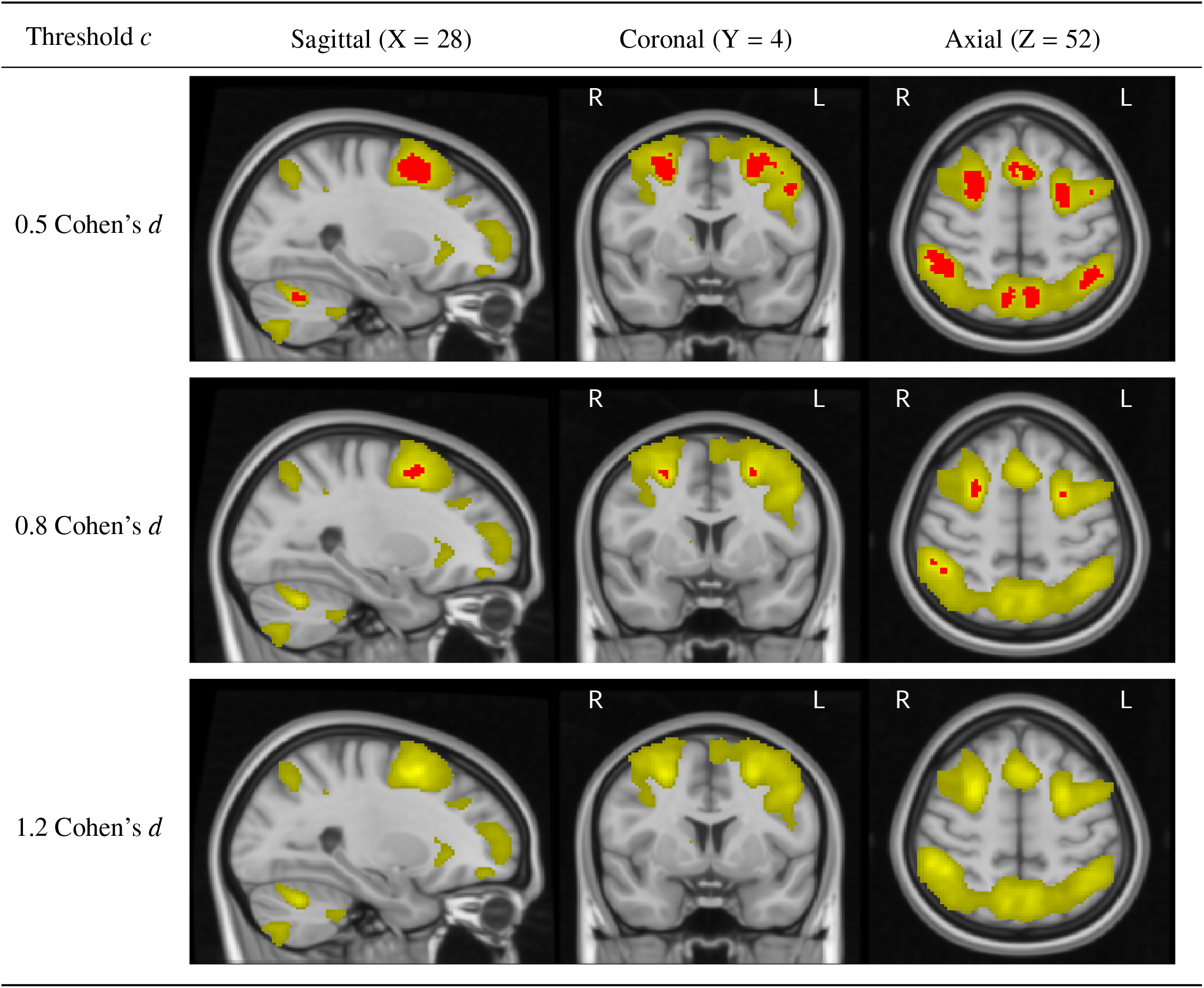
Comparing the upper CSs (red voxels) computed with Algorithm 3. on the HCP working memory task data (same slice views as Fig. 11) with the thresholded *t*-statistic results obtained by applying a traditional group-level one-sample *t*-test, voxelwise *p* < 0.05 FWE correction (green-yellow voxels). While the thresholded statistic map contains a single cluster covering a sizable portion of the parietal lobe across both hemispheres (axial slices), the upper CSs have localized the precise areas of the precuneus and anglur gyrus where we can confidently declare a Cohen’s *d* effect size of at least 0.5. This demonstrates how the CSs can provide improved spatial specificity in determining regions with practically significant activation.

## 5. Discussion

### 5.1 Spatial Inference on Cohen’s d Effect Size

To fully appreciate the outcomes of a neuroimaging study, information about the magnitude (as well as presence) of effects must be reported at the end of an investigation. It is only with this knowledge that one can truly determine the practical relevance (and potential clinical importance) of any discoveries made during the analysis. In this work, we have presented three methods to create Confidence Sets for Cohen’s *d* effect size maps, providing formal confidence statements on regions of the brain where the Cohen’s *d* effect size has exceeded a specified activation threshold, alongside regions where the effect size has not surpassed this threshold. Both of these statements are made simultaneously across the entire brain, enabling researchers to pinpoint the precise regions where meaningful differences have occured as well as identifying areas that have not responded to the task. This is in contrast to the traditional statistical approach, where a test statistic (e.g. a t-statistic) or a p-value can only quantify the compatibility between the observed data and what would be expected under the null-hypothesis of no activation, and no information is provided about the effect size when a finding is deemed to be statistically significant. Since the test statistic is confounded by the sample size of the study, it is not possible to implement a similar framework to obtain Confidence Sets for statistic images; as sample sizes increase test statistics also become arbitrarily large, until ultimately there is enough statistical power to declare even the smallest of effects as statistically significant. On the other hand, for larger samples we expect the red upper CSs to converge towards the population parameter 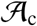 representing the set of voxels with a true population effect magnitude above a purposeful activation threshold c. At the same, while the logic of null-hypothesis testing can never lead to the acceptance of the null hypothesis, the blue lower CSs allow users to make inference on where effect sizes have not attained a sufficient activation threshold.

The use of CSs for inference on effect size may also help to alleviate issues associated with hypothesis testing for studies with lower statistical power. Specifically, for studies with small sample sizes it has been reported that applying traditional inference procedures can lead to spurious or irreproducible results with considerably inflated observed effect sizes (Poldrack et al., 2017, Cremers et al., 2017). In regards to the latter, this is often caused by a form of selection bias known as the ‘winners curse’ (Reddan et al., 2017, Button et al., 2013), whereby voxels whose observed effect size has exceeded their expected performance are intrinsically more likely to be determined as statistically significant. This becomes a problem as magnitudes are commonly reported *only* for significant voxels, a practice that leads to positively biased effect estimates. Our analysis results from the Human Connectome Project dataset exemplify how the CSs can help to resolve this issue. In Fig. 11, the yellow point estimate ‘best guess from the data’ clusters identified a number of voxels with a Cohen’s *d* effect size greater than 1.2 that were also included in the thresholded statistic map obtained from applying a one-sample *t*-test, voxelwise *p* < 0.05 FWE correction (Fig. 12). However, by synthesizing information about the effect magnitude as well as the *reliability* of the estimate, the CSs presented in Fig. 11 affirm that there are in fact no voxels that can be confidently declared to have an effect size larger than 1.2. On the contrary, only a handful of brain regions were contained in the red upper CS asserting a Cohen’s *d* effect size exceeding 0.5 for the HCP working memory task data.

In our previous effort we described a method to obtain CSs for unstandardized percentage BOLD change maps, rather than the Cohen’s *d* images that have served as our main focus here. The use of Cohen’s *d* instead of %BOLD is likely to be advantageous due to complications associated with the BOLD effect. As discussed in our previous work, %BOLD effect sizes have been shown to modulate according to acquisition parameters such as the scanner field strength or MRI pulse sequence, and inhomogenieties in vascularity between different brain regions can cause further variation in the BOLD response. At a more rudimentary level, there are also difficulties involved in obtaining percentage BOLD change images. While all of the main fMRI software packages provide contrast of parameter estimate maps, each of the three most widely-used analysis packages (AFNI, FSL & SPM) scale the raw data differently; the parameter estimates are often given in arbitrary units which deviate between packages. Conversion to percentage BOLD change therefore requires a software-dependent normalization, where one must take into consideration how to appropriately scale the data, design matrix and analysis contrasts. While this can be cumbersome and prone to human error, conversion to Cohen’s *d* is relatively simple. Due to the straightforward relationship between the Cohen’s *d* effect size and the one-sample t-statistic 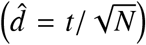, users can easily generate Cohen’s *d* images from the unthresholded *t*-statistic maps created by all the main neuroimaging packages. For all of these reasons, the Cohen’s *d* CS maps may be more suitable for comparison between studies.

In this work, we have used classifications of the Cohen’s *d* effect size as initially suggested in Cohen (2013), describing 0.5 as a ‘medium’ effect, 0.8 as ‘large’, and 1.2 as ‘very large’. While these benchmarks provide basic descriptors of effect size, in general we recommend that users take appropriate steps to contextualize what sort of magnitude constitutes a meaningful finding in their own study. Users should factor in the aims of their investigation, the quality of the study and, if possible, the effect sizes reported in similar previous efforts before choosing a threshold. Obtaining the CSs for the Human Connectome Project contrast data in this work was computationally quick, with each analysis taking less than one minute for all three proposed algorithms. Therefore, one possible strategy is to evaluate a variety of different *c*’s on pilot or historical data before fixing a value to use on a study of interest.

While we have developed the Cohen’s *d* CSs for a one-sample model, the methods presented here may also be applied to the general linear model ***Y***(***s***) = ***Xβ***(***s***) + ***ϵ***(***s***), where ***X*** is the design matrix and ***β***(***s***) is the vector of unknown coefficients. In this setting, for a contrast vector ***w***, the quantity of interest would be the standardized contrast ***w***^*T*^***β***(***s***)/*σ*(***s***) (instead of the Cohen’s *d* effect size *μ*(***s***)/*σ*(***s***)). The method would be carried out similarly, except, for example, the normalized standard deviation of the contrast estimate 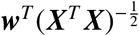 would need to be considered in the construction of the CSs (replacing the 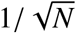 term in the one-sample CS constructions for the three algorithms). It is important to note that the mathematical results underpinning this work in Telschow, Davenport, and Schwartzman (2020) have as yet only been provided for the one-sample model, which is why this model has been our primary focus here. Nevertheless, a further intuition on applying the method to the general linear model can be obtained from *BTSN*, where we developed the CSs in the general linear model setting for raw effect size images.

### 5.2. Three Algorithms for Cohen’s d Confidence Sets

In this work, we have theoretically motivated three algorithms for obtaining Cohen’s *d* CSs. Our simulation results in Sections 4.1 and 4.2 have demonstrated differences in the coverage performance for each of these algorithms. Across all sets of simulation results, empirical coverage for Algorithm 1. came above the nominal level, with particularly severe over-coverage for 3D simulations carried out on large synthetic signals when small sample sizes were used (Figs. 7, 8 and 9). The cause for such poor performance here is likely to be due to the variance term used to construct the CSs in Algorithm 1. Recalling the derivations in Section 2.2, the variance term 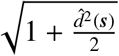 used for Algorithm 1. was chosen as an estimator of the variance of the limiting Gaussian field 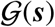. Therefore, while we expect this term to correctly approximate the variance of the bootstrap approximating field asymptotically, our theory provides no indication about the accuracy of this term in small samples. The over-coverage seen in our simulation results suggests that this term overestimates the true variance of the approximating field when the sample size is low. While there was some improvement in Algorithm 1.’s results as *N* increased, even for the largest sample size we analyzed, *N* = 480, empirical coverage for the other two methods was consistently closer to the nominal target level.

Algorithm 2. and Algorithm 3. performed well in all of our 2D and 3D simulations. For simulations using a 95% confidence level, the empirical coverage performance of these two methods was remarkably similar for *N* ≥ 120 (in most cases, slightly above the nominal target). It is therefore difficult to conclude which method should be implemented in practice. For our 3D simulations (Figs. 7 to 10), Algorithm 2.‘s empirical coverage results fell slightly closer to the nominal level in most cases. However, the results for Algorithm 3. were more robust to changes in sample size; for the UK Biobank and multiple spheres simulations (Figs. 9 and 10), the empirical coverage performance for Algorithm 3. was relatively uniform in terms of sample size, while Algorithm 2.’s coverage fell below the nominal level for *N* = 60. This may imply a further drop-off in performance for Algorithm 2. on data with sample sizes below 60, suggesting that Algorithm 3. could be favourable here.

From a theoretical standpoint, the variance-stabilizing transformation approach used in Algorithm 3. assumes that the observations are Gaussian, while this is somewhat relaxed for Algorithm 2., where the bootstrap is applied to estimate the standard deviation directly from the data. While this supports that Algorithm 2. may be preferable for non-Gaussian data, in our Human Connectome Project analyses the CSs maps obtained using both methods (Fig. 11 for Algorithm 3., Fig. S.2 for Algorithm 2.) were virtually identical, indicating that both methods could be equally effective for fMRI data with sample sizes on the order of the HCP.

It is noticeable that the asymptotic coverage for all three procedures appeared to converge to above the nominal level in nearly all of our simulations. As well as this, the size of the overcoverage varied depending on the signal type and the confidence level (in all cases, the degree of over-coverage was higher for the smaller confidence level of 1 – *α* = 0.80 compared to the larger confidence level of 1 – *α* = 0.95). We do not believe this is due to changes in the signal type per se, but instead due to inaccuracies in the interpolation method used to assess if coverage was obtained (i.e. if 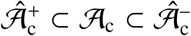) caused by the resolution of the lattice, positively biasing results for signals with a longer boundary 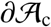. The assessment method evaluates whether coverage holds at a discrete set of sub-sampled points on 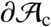, but as the boundary length becomes longer, this set of discrete points becomes relatively less dense within the true, continuous boundary. Violations of the subset condition are therefore more likely to be missed for signals with a longer boundary, which could explain why, for example, the empirical coverage results for the large spherical effect were systematically higher than for the small spherical effect(Figs. 7 and 8). This line of reasoning is also consistent with the greater levels of over-coverage seen for the lower confidence levels, as more violations of coverage should have occurred here, but this also meant there was a higher chance that violations could be missed. We discussed this issue in further detail in Section 5.2 of *BTSN*.

## 6. Data Availability

We have used data from The Human Connectome Project and UK Biobank. All code used for the simulations and analysis of HCP data are available at: https://github.com/AlexBowring/Confidence_Sets_Manuscript.

## 7. Acknowledgements

A.B. was supported by an Early Career Research Fellowship through the Nuffield Department of Population Health. T.E.N. was supported by the Wellcome Trust (100309/Z/12/Z). F.T. and A.S. were partially supported by NIH grant R01EB026859.

Data were provided [in part] by the Human Connectome Project, WU-Minn Consortium (Principal Investigators: David Van Essen and Kamil Ugurbil; 1U54MH091657) funded by the 16 NIH Institutes and Centers that support the NIH Blueprint for Neuroscience Research; and by the McDonnell Center for Systems Neuroscience at Washington University.

## Appendix A. Variance-Stabilizing Transformation of Cohen’s *d* Estimator

### Theorem 1.

*Assume the one-sample Gaussian model described in Section 2.2. For fixed N, let:*

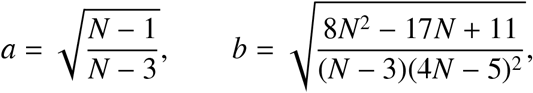

and define

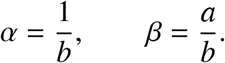

*Then for* 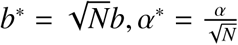, and 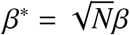, *define the transformation ζ*: ℝ → ℝ *as*:

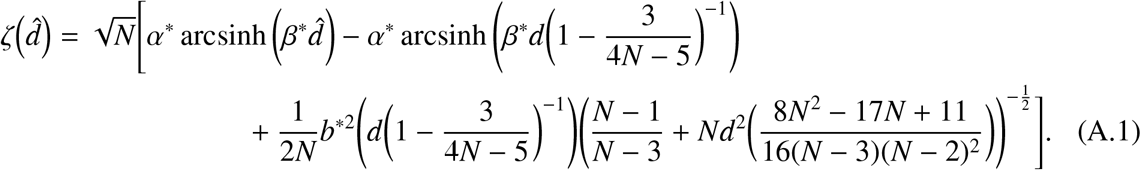

*Then the random variable* 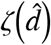 *has, approximately, zero mean and unit variance*.

*Proof.* We closely followthe workings given in the ‘**2. Noncentral t.**’section of Laubscher (1960). We have shown that 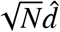 is distributed by a noncentral *t*-distribution with noncentrality parameter 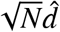 and *N* – 1 degrees of freedom. Defining:

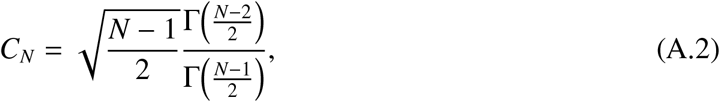

where Γ is the gamma function, then the expectation and variance of 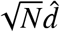 are:

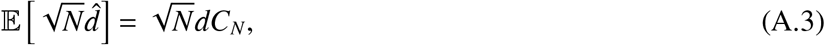

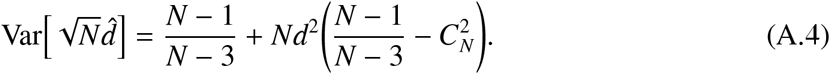

It is known that *C_N_* is well-approximated by the polynomial

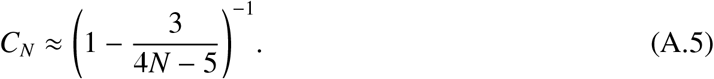

Substituting into equations (A.3) and (A.4), we deduce approximations of the expectation and variance of 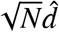,

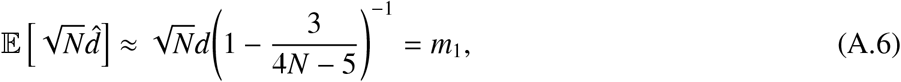

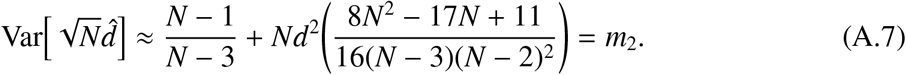

Now, noting that:

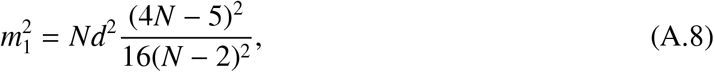

then *m*_2_ can be expressed in the form

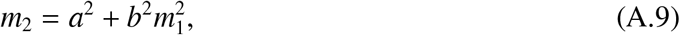

where 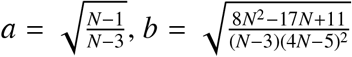. Using the variance expression in (A.9) and applying Corollary 1. in Laubscher, the approximate variance-stabilizing transformation of 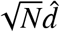 is given by

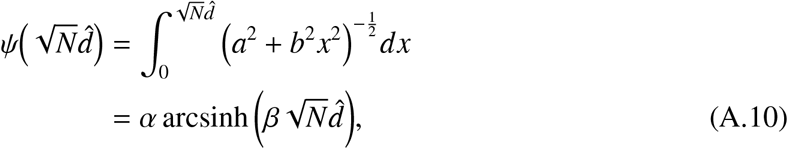

where 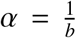 and 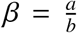. The quadratic Taylor approximation of 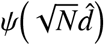 about the point *m*_1_ is given by

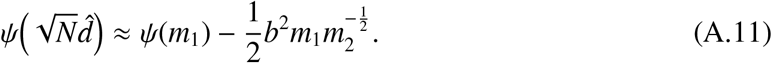

Therefore, the random variable:

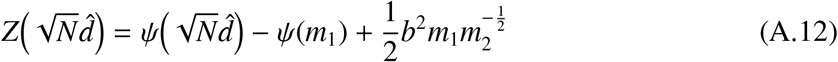

will have, approximately, mean zero and unit variance. Substituting the precise expressions for *ψ,m*_1_, and *m*_2_ into (A.12) yields

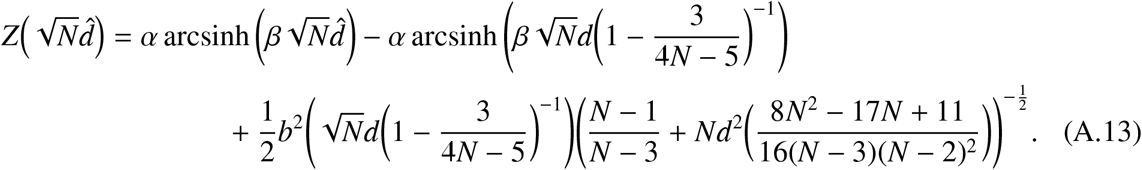

At this point, we have established a variance-stabilizing transformation in terms of 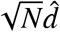, when in practice, we require a transformation in terms of the Cohen’s *d* estimator 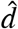. This is possible by applying a change of variables to *b, α* and *β*. Defining 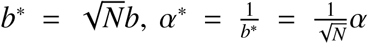, and 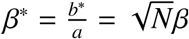, substituting into (A.13) obtains the desired transformation:

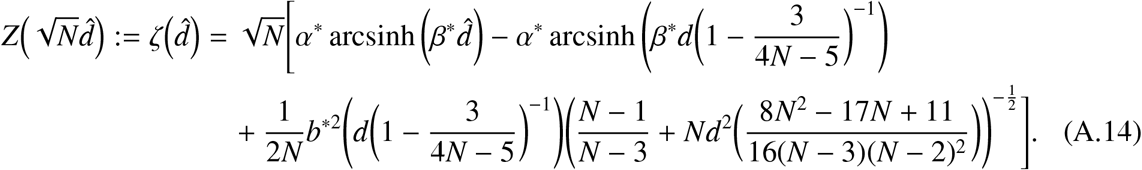

## Appendix B. Supplementary Human Connectome Project Results

**Figure S.1:**
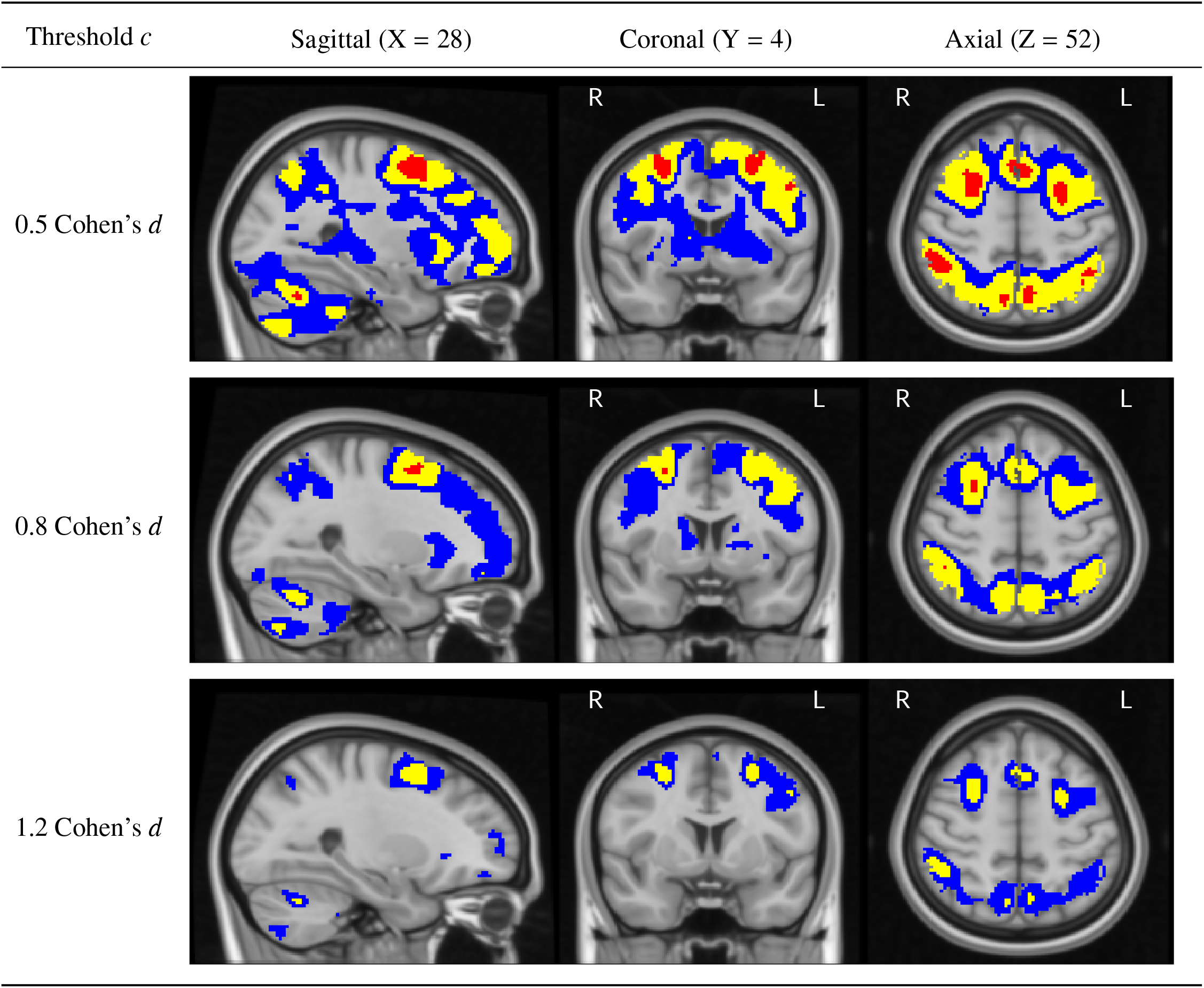
Slices views of the Cohen’s *d* Confidence Sets obtained from applying Algorithm 1. to the HCP working memory task data, using three Cohen’s *d* effect size thresholds, *c* = 0.5, 0.8 and 1.2. Comparing with Fig. 11 and Fig. S.2, the CSs presented here are slightly more conservative than the corresponding CSs obtained with Algorithm 2. and Algorithm 3. (in the sense that the red upper CSs here are smaller, and blue lower CSs are larger). This is consistent with the simulation results obtained in Section 4.1 and 4.2, where the empirical coverage for Algorithm 1. was consistently larger than the other two methods.

**Figure S.2:**
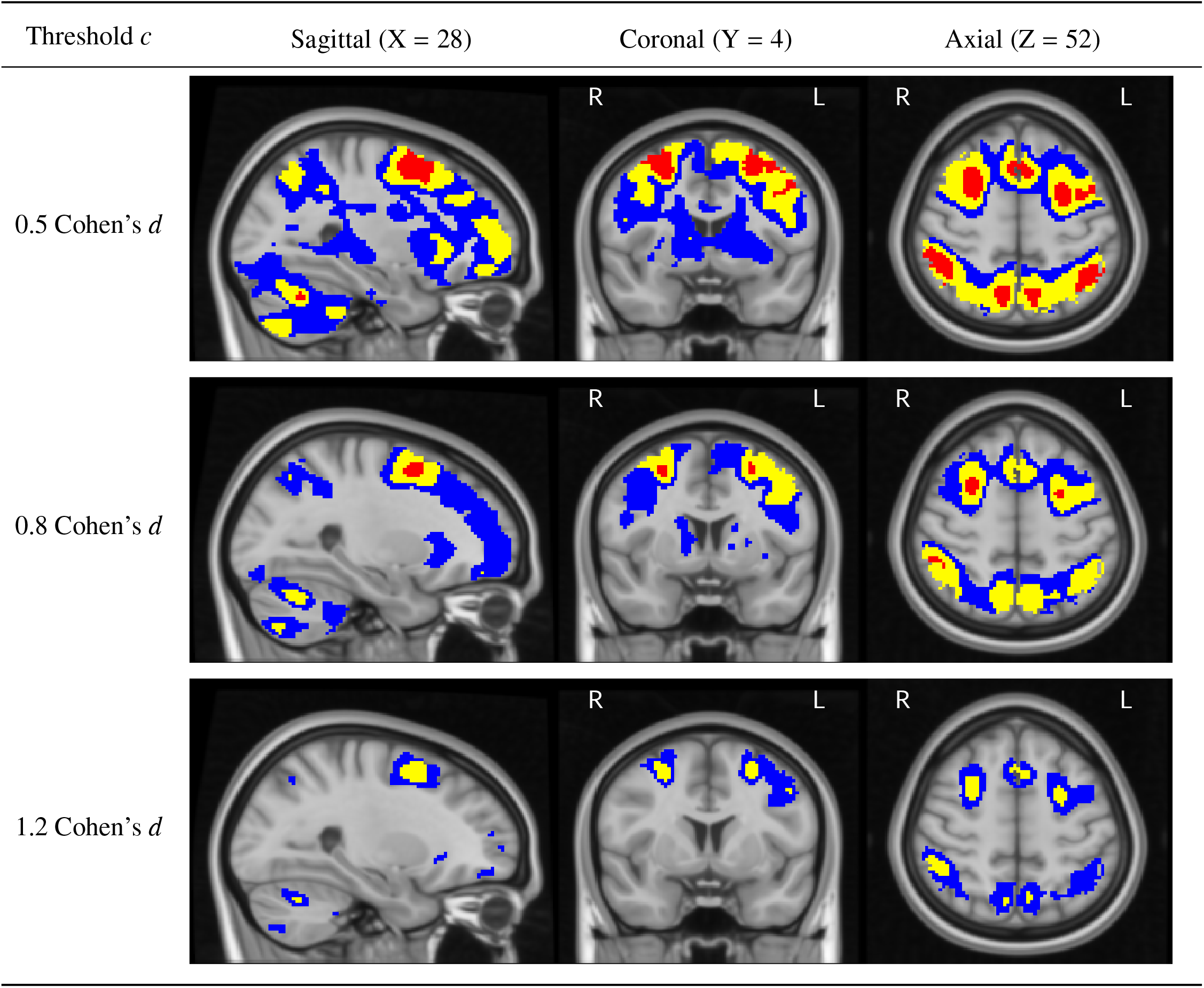
Slices views of the Cohen’s *d* Confidence Sets obtained from applying Algorithm 2. to the HCP working memory task data, using three Cohen’s *d* effect size thresholds, *c* = 0.5, 0.8 and 1.2. Comparing with Fig. 11, the upper and lower CSs presented here are almost identical to the corresponding CSs obtained with Algorithm 3.

## References

Fidel Alfaro-Almagro, Mark Jenkinson, Neal K Bangerter, Jesper L R Andersson, Ludovica Griffanti, Gwenaëlle Douaud, Stamatios N Sotiropoulos, Saad Jbabdi, Moises Hernandez-Fernandez, Emmanuel Vallee, Diego Vidaurre, Matthew Webster, Paul McCarthy, Christopher Rorden, Alessandro Daducci, Daniel C Alexander, Hui Zhang, Iulius Dragonu, Paul M Matthews, Karla L Miller, and Stephen M Smith. Image processing and quality control for the first 10,000 brain imaging datasets from UK biobank. Neuroimage, 166:400–424, February 2018.

Alexander Bowring, Fabian Telschow, Armin Schwartzman, and Thomas E. Nichols. Spatial confidence sets for raw effect size images. NeuroImage, 203:116187, 2019. ISSN 1053-8119. doi: https://doi.org/10.1016/j.neuroimage. 2019.116187. URL http://www.sciencedirect.com/science/article/pii/S1053811919307785.

Katherine S Button, John P A Ioannidis, Claire Mokrysz, Brian A Nosek, Jonathan Flint, Emma S J Robinson, and Marcus R Munafo. Power failure: why small sample size undermines the reliability of neuroscience. Nat. Rev. Neurosci., 14(5):365–376, May 2013.

John T Cacioppo, Stephanie Cacioppo, Gian C Gonzaga, Elizabeth L Ogburn, and Tyler J VanderWeele. Marital satisfaction and break-ups differ across on-line and off-line meeting venues. Proc. Natl. Acad. Sci. U. S. A., 110 (25):10135–10140, June 2013.

Joshua Carp. The secret lives of experiments: methods reporting in the fMRI literature. Neuroimage, 63(1):289–300, October 2012.

Jacob Cohen. Statistical power analysis for the behavioral sciences. Routledge, 2013.

Henk R Cremers, Tor D Wager, and Tal Yarkoni. The relation between statistical power and inference in fMRI. PLoS One, 12(11):e0184923, November 2017.

Matthew F Glasser David C. Van Essen. The human connectome project: Progress and prospects. Cerebrum, 2016, 2016.

Matthew F Glasser, Stamatios N Sotiropoulos, J Anthony Wilson, Timothy S Coalson, Bruce Fischl, JesperL Andersson, Junqian Xu, Saad Jbabdi, Matthew Webster, Jonathan R Polimeni, David C Van Essen, Mark Jenkinson, and WU-Minn HCP Consortium. The minimal preprocessing pipelines for the human connectome project. Neuroimage, 80:105–124, October 2013.

Javier Gonzalez-Castillo, Ziad S Saad, Daniel A Handwerker, Souheil J Inati, Noah Brenowitz, and Peter A Bandettini. Whole-brain, time-locked activation with simple tasks revealed using massive averaging and model-free analysis. Proc. Natl. Acad. Sci. U. S. A., 109(14):5487–5492, April 2012.

Ahmad R Hariri, Alessandro Tessitore, Venkata S Mattay, Francesco Fera, and Daniel R Weinberger. The amygdala response to emotional stimuli: A comparison of faces and scenes. Neuroimage, 17(1):317–323, September 2002.

Nico F Laubscher. Normalizing the noncentral t and F distributions, 1960.

Paul E Meehl. Theory-Testing in psychology and physics: A methodological paradox. Philos. Sci., 34(2):103–115, June 1967.

Karla L Miller, Fidel Alfaro-Almagro, Neal K Bangerter, David L Thomas, Essa Yacoub, Junqian Xu, Andreas J Bartsch, Saad Jbabdi, Stamatios N Sotiropoulos, Jesper L R Andersson, Ludovica Griffanti, Gwenaëlle Douaud, Thomas W Okell, Peter Weale, Iulius Dragonu, Steve Garratt, Sarah Hudson, Rory Collins, Mark Jenkinson, Paul M Matthews, and Stephen M Smith. Multimodal population brain imaging in the UK biobank prospective epidemiological study. Nat. Neurosci., 19(11):1523–1536, November 2016.

Regina Nuzzo. Online daters do better in the marriage stakes, 2013.

Regina Nuzzo. Scientific method: statistical errors. Nature, 506(7487):150–152, February 2014.

Russell A Poldrack, Chris I Baker, Joke Durnez, Krzysztof J Gorgolewski, Paul M Matthews, Marcus R Munafo, Thomas E Nichols, Jean-Baptiste Poline, Edward Vul, and Tal Yarkoni. Scanning the horizon: towards transparent and reproducible neuroimaging research. Nat. Rev. Neurosci., 18(2):115–126, February 2017.

Marianne C Reddan, Martin A Lindquist, and Tor D Wager. Effect size estimation in neuroimaging, 2017.

Michael J Rosenfeld, Reuben J Thomas, and Sonia Hausen. Disintermediating your friends: How online dating in the united states displaces other ways of meeting. Proc. Natl. Acad. Sci. U. S. A., 116(36):17753–17758, September 2019.

William W Rozeboom. The fallacy of the null-hypothesis significance test. Psychol. Bull., 57(5):416–428, 1960.

Max Sommerfeld, Stephan Sain, and Armin Schwartzman. Confidence regions for spatial excursion sets from repeated random field observations, with an application to climate. J. Am. Stat. Assoc., 113(523):1327–1340, July 2018.

Fabian J. E. Telschow, Samuel Davenport, and Armin Schwartzman. Functional delta residuals and applications to functional effect sizes, 2020.

Choong-Wan Woo, Anjali Krishnan, and Tor D Wager. Cluster-extent based thresholding in fMRI analyses: pitfalls and recommendations. Neuroimage, 91:412–419, May 2014.

